# Matrilysin (MMP-7) Restricts RSV Infection and Airway Inflammation through Proteolysis of CCR3 and RSV-G

**DOI:** 10.64898/2026.09.02.748627

**Authors:** Ihssan Kdah, Hassan Ait Benhassou, Salaheddine Redouane, Vincent Wellemans, Houda Houari, Remi Meunier, Richard Le Naour, Meriem Khyatti, Abdelilah S. Gounni, Bouchaib Lamkhioued

## Abstract

Airway epithelium forms a frontline barrier against microbial invasion through mucociliary clearance and the secretion of antimicrobial and immunomodulatory mediators. Matrix metalloproteinases (MMPs), traditionally viewed as tissue-remodelling enzymes, are also implicated in inflammatory processes and are recognized for their capacity to shap host–pathogen interactions. This study investigates the role of MMP-7 (matrilysin) in airway inflammation and the pathogenesis of respiratory syncytial virus (RSV) infection. In silico analyses predicted that MMP-7 binds CCR3, the receptor for eotaxin-1/CCL11, and that it also engages the region of the RSV G protein that docks onto CCR3, pointing to a shared biochemical mechanism interfering with both chemokine binding and viral attachment. Mechanistically, MMP-7 proteolytically cleaved CCR3, disrupting eotaxin-1/CCL11 binding and reducing eosinophil activation and chemotaxis, whereas CX3CR1 was unaffected. Importantly, biophysical measurements confirmed that MMP-7 binds and cleaves the RSV G protein, consequently restricting RSV entry and replication in airway epithelial cells. In vivo, MMP-7-deficient mice displayed higher RSV infection, exacerbated airway inflammation and increased mucus production following RSV challenge. Collectively, these findings identify MMP-7 as a key epithelial antiviral effector coordinating both antiviral defense and the regulation of pulmonary inflammation.

## INTRODUCTION

Respiratory syncytial virus (RSV) is the leading cause of lower respiratory tract infection (LRTI) in infants and a major cause of morbidity in older adults^1^. Following a marked disruption of its seasonality, RSV has resurged worldwide, with out-of-season epidemics reported in several regions. RSV has evolved sophisticated strategies to exploit host cellular machinery, enabling efficient replication and dissemination^2,3^. Successful RSV infection hinges on the direct interaction of the RSV-G attachment protein with cell-surface receptors including CX3CR1, through a CX3C motif within its central conserved domain and, more recently proposed, CCR3^4,5^. This binding event triggers conformational changes in the fusion (F) glycoprotein and facilitating viral and cell membrane fusion and viral entry. Beyond their role in attachment, RSV proteins also modulate host matrix metalloproteinase (MMP) activity, and MMPs have in turn been implicated in airway inflammation and remodelling during infection. Emerging evidence suggests that MMPs shape interactions between respiratory viruses and their host-cell receptors, yet whether a specific MMP acts directly at this interface remains unknown for RSV.

Airway epithelium represents the initial site of host-pathogen interaction, driving the release of inflammatory cytokines and chemokines that shape the early immune response both before and during infection. A key aspect of this host innate immune response to viral respiratory infections involves also the activation and regulation of proteases, including MMPs: a family of zinc-dependent endopeptidases that have emerged as significant contributors to the pathogenesis of viral pneumonia^6^. During respiratory infections, the airway epithelium responds by increasing its defences and producing mediators such as cytokines, chemokines, and MMPs which lead to airway inflammation and cell accumulation^7^. This inflammatory response includes the recruitment of neutrophils, which usually serve to remove microorganisms that are not cleared by the epithelium itself^8,9^. Both epithelial cells and activated neutrophils release MMPs and Neutrophil Serine Proteases (NSPs) that target ECM proteins, thereby shaping the inflammatory response and lung injury^10^. However, it is increasingly clear that matrix degradation is neither the sole nor the common mission of these enzymes, as they can also control chemokine activity. This control can be direct, by MMP-mediated cleavage of these molecules, which results in enhancement, inactivation or antagonism of chemokine activities^11^. It can be indirect, by the proteinases acting on other substrates that bind, retain or concentrate the chemotactic molecules in particular locations^11^.

A key regulatory function of Matrix Metalloproteinases (MMPs) involves the proteolytic conversion of chemokines from potent chemoattractants into antagonistic derivatives, thereby attenuating the recruitment of inflammatory cells to sites of injury. This repressive processing is exemplified by MMP-2, which cleaves the four amino (N)-terminal residues of monocyte chemotactic protein-3 (MCP-3/CCL7) ^12^ and stromal cell-derived factor-1 (SDF-1/CXCL12) ^13^. While these truncated isoforms retain the ability to bind their cognate receptors (CCR1 and CXCR4, respectively), they lose their chemotactic efficacy and instead function as competitive antagonists. Similarly, MMP-1, -3, -13, and -14 have been shown to cleave the N-termini of MCP-1, MCP-2, and MCP-4, generating antagonist factors that further suppress the inflammatory cascade ^14^. The eosinophil chemoattractant eotaxin-1/CCL11 is also cleaved at the N terminus by Dipeptidyl peptidase IV (CD26), which results in a loss of chemotactic potency, while retaining the ability of the chemokine to bind to CCR3^15^. Although MMPs have long been considered to augment inflammation and the associated tissue damage, these observations further imply that they can also dampen inflammatory processes. By contrast, the processing of IL-8 by MMP-9 markedly increases its chemotactic activity^16^. The bioavailability of chemokines is also regulated by their immobilization to the ECM or cell surfaces^17^. *In vivo*, chemokines form chemotactic gradients by binding to accessory macromolecules (typically the glycosaminoglycan side chains of proteoglycans), thereby providing directional cues to migrating leukocytes^18,19^. These studies indicate that by acting on these accessory molecules, MMPs can indirectly regulate chemokine activity and, in turn, the influx of inflammatory cells. MMPs may also possess antiviral activity and play a role in regulating lung inflammation. Indeed, emerging evidence suggests these enzymes facilitate or inhibit viral entry, promote inflammation, and contribute to tissue damage during the pathogenesis of SARS-CoV-2 and RSV infections^20,21–24^.

Beyond the physical interaction between RSV-G and host receptors, RSV relies on activation of its attachment protein (G) and fusion protein (F) by host proteases, underscoring the importance of cell-surface and intracellular proteases in the infectious cycles of this respiratory pathogen. Precisely, it has been demonstrated that the cleavage of the RSV-F protein is required for its surface expression and the fusion of the viral envelope with the target cell membrane ^25^. Moreover, MMP-12^-/-^ mice infected with RSV displayed increased morbidity and significantly higher viral loads^26^. This was accompanied by reduced IFN-levels, despite strong IL-1 and IFN-responses observed in vivo. Additionally, the study revealed that MMP-12 is translocated to the cell nucleus and acts as a transcription factor, thereby amplifying the antiviral immune response^26^. Recent studies demonstrate that MMP-9 plays a critical role during pulmonary viral infection. MMP-9 expression is required for neutrophil recruitment to the lung, cytokine production, viral clearance, and is associated with reduced airway hyperresponsiveness (AHR) following RSV infection^27^. However, in MMP-9□/□mice, protection from influenza A virus (IAV)–associated mortality is associated with reduced lung tissue damage and enhanced adaptive immune responses, leading to more efficient viral clearance^28^. SARS-CoV-2, another respiratory virus, similarly requires proteolytic activation by host proteases for cell entry. During virion maturation, furin cleaves the S protein at the S1/S2 junction, priming it for later processing ^29^. Upon ACE2 engagement, TMPRSS2 cleaves the S2′ site, enabling membrane fusion ^30,31^. Alternatively, in cells with low TMPRSS2 expression, cathepsin L activates the S protein within endolysosomes, enabling entry via the endocytic pathway ^32^ ^33^. These coordinated proteolytic events are critical determinants of viral tropism and pathogenesis, as observed in other respiratory viruses such as influenza^34^. Understanding the potential roles of MMPs in RSV infection may thus provide valuable insights into the mechanisms of viral pneumonia and support the development of targeted therapeutic strategies.

In this report, we provide evidence that CCR3, the receptor for eotaxin-1/CCL11 which plays a central role in allergic diseases and RSV infection^5^, is a novel substrate for MMP-7. We demonstrate that MMP-7 cleaves CCR3, thereby inhibiting the activity of eotaxin/CCL11, a chemokine that recruits eosinophils. Moreover, we show that CCR3 cleavage reduces RSV infection in airway epithelial cells. Beyond its action on CCR3, MMP-7 is also capable of cleaving RSV G attachment glycoprotein resulting in decreased RSV infection of host cells. Collectively, these findings suggest a novel mechanism by which MMPs, particularly MMP-7, may contribute to the control of airway infections and inflammatory conditions associated with tissue eosinophilia.

## MATERIALS AND METHODS

### Materials

Biotin-conjugated recombinant human chemokines (eotaxin-1/CCL11, MIP-1α/CCL3 and MIP-1β/CCL4) were obtained from R&D Systems (Minneapolis, MN). Goat anti-human CCR3 antibody was purchased from Santa-Cruz Biotechnology (California, USA). For flow cytometry analysis, anti-CCR3-FITC-conjugated Ab and anti-CCR3-PE-conjugated Ab were from R&D system (Minneapolis, MN). Anti-CD16 and anti-CD3 Abs were obtained from Miltenyi Biotec (Aubrun, USA). Metalloproteinases (MMP-1, -2, -3, -7, -9, -10, -12, -13) were obtained from Biomol International (PA – USA). Unless stated otherwise, all other reagents were obtained from Sigma-Aldrich.

### Cells and culture conditions

CCR3^+^ cells from two sources were used. Ghost cell transfectants are human osteosarcoma cell lines (HOS) stably transfected with different chemokine receptors were obtained through the AIDS Research and Reference Reagent Program, Division of AIDS, NIAID, NIH, USA: Ghost.CCR1, Ghost.CCR3, Ghost.CCR5, Ghost.CXCR4 and Ghost.CX3CR1 from Dr. Vineet N. Kewal Ramani and Dr. Dan R. Littman^35^. Cell cultures were maintained at 37°C in Dulbecco’s modified Eagle’s medium (Invitrogen) supplemented with 10% heat-inactivated fetal calf serum (FCS), Glutamax, antibiotics (100 µg of hygromycin/ml, 500 µg of G418/ml, and 1 µg of puromycin/ml)^5^. The medium was replaced every 2–3 days, and cells were passaged with 0.5% trypsin and 1mM EDTA once confluence was reached. Cell Fresh medium was replaced every 2 days. The second source of cells was human peripheral blood eosinophils that were isolated by negative selection using immunomagnetic beads, as previously described^5^. Briefly, mononuclear cells and granulocytes were purified from peripheral blood by Ficoll-Paque (Pharmacia) density centrifugation. Granulocytes were obtained by dextran sedimentation. Human eosinophils were further purified by negative selection with anti-CD16 and anti-CD3-coated immunomagnetic microbeads using a Magnetic Cell Sorting system (MACS; Miltenyi Biotec Inc., Sunnyvale, CA) at 4°C. The degree of eosinophil purity, estimated after Giemsa staining, ranged from 92 to 100%.

Sf9 insect cells were obtained from BD Bioscience and were grown in TNM-FH insect cell medium (BD Bioscience) containing 10% fetal bovine serum (FBS) (Life Technologies).

### Structure prediction

For 3D structure modeling, the Uniprot database was used to obtain the FASTA sequences of MMP-7 and MMP-9, along with their potential partners CCR3 and the RSV G protein. ColabFold, a condensed version of AlphaFold tailored for Google Colab, was used to do structural prediction for the docking of different complexes: MMP-7–CCR3, MMP-7–G protein, MMP-9–CCR3 and MMP-9– G protein. To improve prediction accuracy, multiple sequence alignments were produced after the FASTA sequences were loaded. Subsequently, the projected Aligned Error matrix, which assesses inter-domain interactions and chain alignment, and the pLDDT score, scaled from 0 to 100 and reflecting confidence in the projected structures, were used to forecast and evaluate the structures’ dependability. ColabFold generates five structural models, each with a confidence value based on prediction accuracy. Several analytical approaches have been used to evaluate the interactions. Specifically, the identification of the conserved regions and the differences between the sequences have been made possible by a sequence coverage card. To distinguish between flexible and uncertain regions, the structures were colored according to the pLDDT score and the various chains. Furthermore, the PAE matrix has enabled estimating the reliability of the generated model and assessing the uncertainty in the alignment of the structures.

### Molecular docking

The predicted docking using AI was validated using a semi-flexible guided approach with HADDOCK 2.4^36^. The 3D structures of the proteins were obtained from the RCSB Protein Data Bank, selecting those with the best resolution and biological relevance, such as active conformation and complete domains. Structures of MMP-7 and MMP-9, along with their potential partners CCR3 and RSV-G protein were used. Before docking, the structures were prepared by removing incomplete chains, irrelevant ligands, ions, and water molecules. Energy minimization was then performed using the Steepest Descent method in GROMACS to achieve a stable, realistic conformation. For each interaction, docking was performed, and models were generated, and the best one was selected based on the HADDOCK score, which is a weighted sum of several energy terms: van der Waals, electrostatic, desolvation, and restraint violation energies, cluster size, Root-mean-square deviation, Buried Surface Area and Z-Score. All complexes generated by docking via Colabfold and HADDOCK were visualised in Chemirax and Pymol.

### Expression and purification of RSV G protein from insect cells

RSV G gene was first cloned into a recombinant baculovirus transfer vector DNA (pACHLT-B, BD Bioscience). Recombinant baculovirus was generated according to the manufacturer’s protocol (BD Bioscience) and a His-fusion protein can be produced by recombinant baculovirus generated using the BD BaculoGoldTM Linearized Baculovirus DNA as previously described^5^. Briefly, DH5α Escherichia coli cells were transfected with pACHLT-B to generate recombinant baculovirus shuttle vector DNA. Sf9 Spodoptera frugiperda cells were then transfected with 2 μg baculovirus shuttle vector DNA with transfection reagents according to the manufacturer’s recommendation (BD Bioscience), and the baculovirus was amplified several times to obtain the optimal viral titer. pVL1392-XylE, a pseudomonas putrida gene XylE cloned into the pVL1392 baculovirus transfer vector, was used as a positive co-transfection control in these experiments. For protein expression, baculovirus stocks were added to Sf9 cells grown in suspension at a density of 1 × 10^6^ cells per ml, and transduced Sf9 cells were collected 2 d after transduction. Recombinant baculoviruses containing the RSV G gene were identified by RT-PCR or western blot analysis of insect cell supernatants and the RSV G recombinant fusion protein was affinity-purified using Ni-NTA agarose beads as recommended by the supplier (BD Bioscience).

### Proteinases treatment of the cells

Human eosinophils and subconfluent CCRs^+^-transfected Ghost-cells (G.CCR^+^) were exposed to MMPs (ranging from 1 to 100 ng/ml) for 4h at 37°C. To assess whether the full-length matrilysin directly cleave CCR3, confluent G.CCR3^+^ cells or eosinophils in 6-well plates were incubated for 4h at 37°C in 2 ml medium without or with recombinant pro-matrilysin activated with 1 mM APMA (RD systems), or MMP catalytic domains. Thereafter, adherent GHOS cells were detached with PBS, 2 mM EDTA for 20 min at 37°C. Cells were washed, counted and used for the western blot analysis, flow cytometry analysis or chemotaxis assays as described bellow. To demonstrate MMP-dependent processing of the chemokine receptors, GM6001 (an MMPs inhibitor) at 1 mM was added to the reaction. In some experiments, A549 cells were stimulated with proinflammatory cytokines (IL-1β and TNF at 25 ng/ml) that are usually used to mimic the inflammatory reaction^5^, and the supernatant was recovered 48h later and added to CCR3-transfected cells in the presence or absence of GM6001 (1 mM) to confirm that the observations are specifically due to MMP enzyme activity.

### Proteinases treatment of viral proteins

Matrix metalloproteinases (MMPs), including active human recombinant MMP-7 expressed in E. coli (Merck, Cat. No. 444270) and recombinant human MMP-2 (BioLegend, Cat. No. 554302), were used to treat recombinant forms of the RSV-G protein (RSV-G protein, His-Tag; Sino Biological, Cat. No. 11070-V08H). Following activation with 4-aminophenylmercuric acetate (APMA), 10 µL of MMP-2 or MMP-7 were each incubated with 10 µL of RSV-G protein at 37 °C overnight. Proteins were separated on 13 % polyacrylamide gel at 4°C and electrophoretically transferred to nitrocellulose membranes (Amersham Pharmacia Biotech) as described below.

### Western blotting

After MMP treatment, cell pellets were incubated in lysis buffer (150 mM NaCl, 10 mM Tris-HCl, pH 7.4, 1 mM EDTA, 1 mM EGTA, 1% Triton X-100, and 0.5% Nonidet P-40) containing a protease inhibitor mixture tablet (Roche Diagnostic Systems). Lysates were incubated at 4°C for 30 min and insoluble debris was removed by centrifugation (14,000 x g for 30 min). Proteins were separated on 13 % tricine-gel at 4°C and electrophoretically transferred to nitrocellulose membranes (Amersham Pharmacia Biotech). Membrane blocking was achieved using 5% non-fat dry milk in TTBS (0.1% Tween 20, 10 mM TBS, pH 7.5) for 2 h at RT and then washing twice for 2 min with TTBS. Membranes were incubated successively with anti-CCR3 Abs (5 μg/ml; Santa Cruz Biotechnology) and alkaline phosphatase-conjugated anti-goat Abs (1/1000, Santa Cruz Biotechnology) for 1h at room temperature. The bound secondary Abs were detected using the CSPD chemiluminescence detection kit (Roche Diagnostic Systems).

For detection of ERK2 phosphorylation, eosinophil protein lysate (10^6^ per sample) was probed using antibodies directed against the dual threonine/tyrosine–phosphorylated MAP kinases, as described previously^37^. SDS-polyacrylamide (10%) gels were used according to the Laemmli protocol. The membranes were incubated with 0.1 μg/mL of primary antibody for 2 hours at room temperature (monoclonal anti-phosphotyrosine antibody, clone 4G10, Upstate Biotechnology, Inc, Lake Placid, NY, or polyclonal anti-ERK2 antibody, monoclonal anti-phospho-ERK2 antibody from Santa Cruz Biotechnology). The membranes were washed 5 times with TBS-T, incubated with horseradish peroxidase–conjungated secondary antibodies (1:10000 dilution of anti-mouse immunoglobulin (Ig) from Santa Cruz Biotechnology) for 20 minutes and washed 5 times with TBS-T. The blots were visualized by the enhanced chemiluminescence system (Amersham) according to the manufacturer’s instructions.

### Cell-binding assays using Flow cytometry analysis

Cells were labelled with a Fluorokine kit for human chemokine receptors (R&D Systems) according to the manufacturer’s instructions. Briefly, adherent CCRS^+^-Ghost cells were first detached from the flask by adding a solution of PBS-EDTA (0.5 M) for 20 min at 37°C. Eosinophils and Ghost cells were washed once with PBS and resuspended at a concentration of 4 ×10^6^ cells/ml. 10 µl of biotinylated recombinant chemokine (eotaxin-1/CCL11, MIP-1α/CCL3 or MIP-1ß/CCL4) reagent was added to 25-µl of the washed cells and incubated for 60 min on ice. As a negative staining control, an identical sample of cells was stained with 10 µl of biotinylated negative control reagent (soybean trypsin inhibitor). Following the incubation period, 10 µl of Avidin-FITC reagent was added, and cells were incubated for an additional 30 min at 4°C in the dark. The specificity of the reaction was assessed by mixing 20 µl of blocking antibody with 10 ml of biotinylated chemokine and incubating for 15 min at RT. Cells were then washed twice, using the buffer provided to remove unreacted Avidin-FITC, resuspended in 200 µl of PBS, and cell-associated immunofluorescence was immediately analysed using FACS machine (BD Biosciences) to determine the level of surface expression of chemokine receptors. CCR3 was also identified using FITC-labelled anti-CCR3 Abs and analysed using FACS machine. Eosinophils were resuspended at a concentration of 1×10^6^ cells/ml, washed once with PBS, and then incubated with purified normal human IgG (Santa Cruz Biotechnology) at 4°C for 20 min to block any non-specific binding. PE-conjugated anti-CCR3 (FAB155P, clone 61828.111; R&D Systems) or control isotype (rat IgG2A; IC006P, clone 54447; R&D Systems) Abs were incubated with the cells at 4°C for 30 min. After three washes with 0.5% PBS-BSA, cells were then resuspended in PBS at 4°C. Cell-associated immunofluorescence was immediately analysed using FACS (BD Biosciences) to determine the level of surface expression of CCR3.

### Chemotaxis assay

Experiments were performed with a 24-well microchemotaxis chamber (NeuroProbe, Cabin John, MD) and conducted as previously described^38^. Migration of G.CCR3 cells and human eosinophils in response to eotaxin-1/CCL11 was performed on a polycarbonate filter (5-µm ad 8 µmm pore size respectively). Briefly, cells were resuspended in RPMI medium supplemented with 0.1% BSA, and 5 ×10^4^ cells were then loaded into the upper chambers and tested for chemoattraction to medium supplemented with either 0.1% BSA (negative control) or increasing doses of eotaxin-1/CCL11. In selected experiments, cells were pre-incubated for 1 hour with 10 µg/mL of anti-CCR3 antibody (MAB155, clone 61828; R&D Systems) prior to chamber loading (data not shown). Migration chambers were then incubated at 37°C in a 5% CO humidified atmosphere for 4 hours. Following incubation, non-migrated cells on the upper surface of the membrane were removed by gentle scraping. Cells that had migrated to the lower surface were stained using an RAL kit (Labonord, France). Migrated cells were quantified by counting five randomly selected high-power fields (×400 magnification) using standard light microscopy^39^.

### Effect of MMPs on RSV-induced syncytium

Hep-2 (human laryngeal carcinoma) and A549 (human type II alveolar epithelial) cell lines were obtained from the American Type Culture Collection (ATCC, Rockville, MD) and maintained in Eagle’s Minimum Essential Medium (EMEM; Gibco BRL, Life Technologies), supplemented with 10% fetal calf serum (FCS), GlutaMAX, 100 U/mL penicillin, and 100 µg/mL streptomycin. Cells were incubated at 37°C in a humidified atmosphere containing 5% CO□. Human respiratory syncytial virus (RSV) strain A2 (Long strain), confirmed to be free of chlamydia and mycoplasma contamination, was obtained from ATCC (Manassas, VA) and propagated in Hep-2 cells as previously described^5,40^.

Viral adsorption was carried out for 1 hour at 37°C, after which 10% EMEM was added. Infected cultures were incubated for 3–5 days until cytopathic effects were observed across the monolayer. Cell culture supernatants were harvested, resuspended in 1-mL aliquots, rapidly frozen using an alcohol/dry ice bath, and stored at –80°C until use. The virus titer of RSV was assessed by quantitative plaque forming assay and maintained titers in the range of 10^6^ to 10^8^ pfu/ml for more than 6 months at -70°C. For the Infectivity inhibition tests twenty-four well tissue culture plates (Falcon, Becton Dickson and Co) were seeded with 3×10^5^ HEp-2 or A549 cells per well in 1 ml of complete medium. After overnight incubation (37°C, 5% CO_2_), triplicate wells were pre-treated for 4 hr at 37°C with recombinant MMP-1, -2, -3, -7, -9, -10, -12, -13 (50 or 100 ng/ml) or heparin (5 µg/ml)^41^. Following removal of MMPs or heparin, the monolayers were infected (1 hr) with RSV at 37°C temperature. Thereafter, the monolayers were washed three times with pre-warmed phosphate buffered saline (PBS) and overlaid with RPMI containing 10% FCS and 0.75% methylcellulose. Three to four days post-infection the cells were fixed and stained with 1% crystal violet. Plaques were counted and percent inhibition of virus infectivity of treated cells was determined versus untreated control wells.

### Virus infection of mice

Pathogen-free 6-week-old BALB/c wild-type (WT) and MMP-7-deficient (MMP-7 /, BALB/c background) mice were obtained from Jackson Laboratory (Bar Harbor, ME, USA) and maintained under specific pathogen-free conditions. All animal procedures were approved by the Institutional Animal Care and Use Committee of the University of Reims and conducted in accordance with the guidelines of the Federation of European Laboratory Animal Science Associations (FELASA) and the European Directive 2010/63/EU on the protection of animals used for scientific purposes. Mice were housed in microisolator cages and provided with sterilized food and water ad libitum. Mice (n = 8 per genotype) were lightly anesthetized and intranasally (i.n.) inoculated with 10 plaque-forming units (PFU) of RSV in 100 μL of sterile PBS. Control animals (n = 8 per genotype) received 100 μL of sterile PBS alone. Mice were monitored daily for clinical signs, and on day 5 post-infection, they were euthanized. Lungs were harvested and processed as described below.

### Lung fixation and histology

The right lung was removed, weighed, resuspended in DMEM and serial dilutions of clarified lung homogenates were plated for viral titer^42^. The right ventricle was flushed with 5 ml clean PBS to flush the left lung vasculature. The trachea was cannulated, and the lungs were inflated with phosphate-buffered formalin (10% formalin) at 20 cm H_2_O pressure. We then removed the lung and submerged it in formalin for fixation and subsequent histological sectioning. Thin sections (10 µM) were cut from paraffin-embedded lungs and stained with hematoxylin and eosin or with Periodic Acid Schiff stain (PAS). As a semiquantitative analysis of mucus induction, goblet cells were counted in PAS-stained tissue sections, counting cells in small to medium airways cut in cross-section. The goblet cell counts were calculated as the average of three airways in lung sections from each mouse and then reported as the mean and standard deviation for each treatment group as previously described^5^.

### Lung Homogenates

The right upper lobe from each mouse was flash-frozen in liquid nitrogen and kept frozen at 80°C. Just before running RSV ELISA assays, the samples were weighed and homogenized in 1 ml of homogenization buffer (Bioo Scientific, Austin, USA) containing one Roche complete protease inhibitor cocktail tablet (Boehringer Mannheim, Germany) and 0.1% Triton-X in 50 ml of phosphate-buffered saline (PBS).

### Statistical analysis

All statistical tests were performed using InStat Software 3.0 (GraphPad Software, San Diego, CA). Data are expressed as mean-G SD of three or more independent experiments with three replicates. Student’s t test (unpaired, two-tailed) was performed when comparing two groups to each other. P values less than 0.05 were considered statistically significant.

## RESULTS

### Docking of CCR3 by MMPs via ColabFold 2

Because airway epithelial cells constitute the first line of defense in the respiratory tract, we anticipated that, following airway infection, these cells might release multiple MMPs that could cleave RSV receptors, thereby modulating viral entry into host cells. Whether a similar mechanism occurs for CCR3—a key receptor for RSV that binds the RSV G protein^5^ —remains unknown. Therefore, we decided to use molecular docking simulations to predict the binding affinity between MMP-7^43^ —a protease secreted by activated airway epithelial cells into the extracellular space^44^—, and CCR-3 (Figure 1). The diagram depicts the computational docking interactions between the CCR3 receptor, and the matrix metalloproteinases MMP-7 and MMP-9, including structural confidence scores (pLDDT), predicted aligned error (PAE) matrices and 3D models of each complex. The predicted local distance difference test (pLDDT) scores ranged from 75.3 to 80.8 across models for MMP-7 and MMP-9, indicating moderate to high confidence in the predicted structures, with consistently higher scores observed for MMP-7. The sequence coverage plots revealed regions of high sequence identity (red) and extensive coverage for MMP-7. In contrast, MMP-9 displayed a more heterogeneous pattern, suggesting the presence of distinct interaction interfaces. The PAE matrices predominantly exhibited blue coloration, indicating low predicted alignment errors and suggesting that the overall predicted structures are reliable. Small red regions at the periphery represent areas of higher uncertainty and generally correspond to flexible segments of the folded protein that are buffered or constrained by surrounding regions. The 3D structures were color-coded according to the pLDDT score, with blue indicating well-resolved folds and yellow/orange highlighting regions of lower confidence, including flexible segments. These colors are not arbitrary; they are mapped directly to the numeric pLDDT values and thus provide a meaningful visualization of model confidence. Regions of lower pLDDT tend to correspond to flexible areas observed in molecular dynamics simulations, reflecting a correlation between model confidence and structural flexibility as discussed elsewhere^45^. Although both MMPs bind to CCR3, MMP-7 exhibited a higher average pLDDT score and more consistent coverage. In contrast, MMP-9, which demonstrates stronger binding in other contexts, showed greater variability in this analysis, potentially reflecting conformational flexibility or differences in epitope recognition. The observed distinctions in interaction profiles suggest generally stable binding for both enzymes, with subtle structural differences.

**Figure 1.**
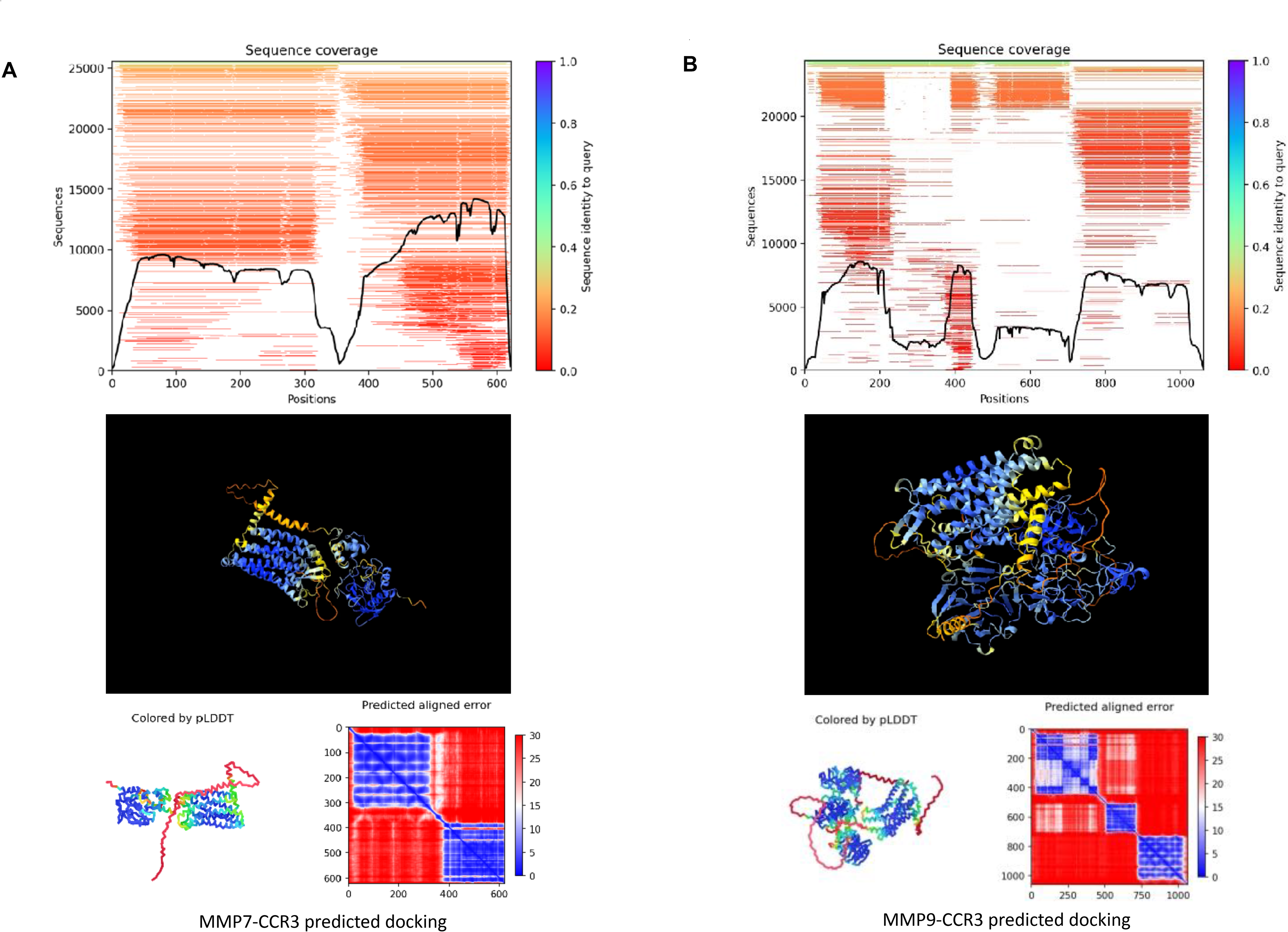
AI-Predicted Docking of CCR3 by MMPs via ColabFold2. 3D models colored by pLDDT scores (blue: high >90; red: low <70) with average confidence shown. PAE plots indicate inter-residue prediction errors (blue: low error; red: high error). MSA coverage plots showing sequence identity (color) and alignment depth (black curve). Low coverage regions correspond to lower model confidence. A: The complex MMP-7-CCR3 (left) shows good overall MSA coverage across the ∼600-residue sequence and relatively reliable predictions (mean pLDDT = 78.729), although a drop in confidence is observed around the central region. B: The second complex MMP-7_CCR3 (right), which is longer (∼1000 residues), exhibits poorly aligned regions in the MSA—particularly around positions 400–600—correlating with locally reduced model confidence (mean pLDDT = 76.093).

### Molecular docking of CCR3 by MMPs via HADDOCK.2

To validate the ColabFold docking predictions of MMP-7/-9 with the CCR3 receptor, we performed a more detailed analysis using HADDOCK platform. This confirms the biologically relevant interactions observed, supporting their potential role in an inflammatory environment following RSV infection (Figure 2). The results indicate a stable interaction between the MMPs and CCR3. Among the clusters generated by HADDOCK, the first cluster of MMP-7-CCR3 complex, provided the informative results, with a ADDOC score of -84.4 ± (0.5), a Z-score of -1.5, and a reasonable cluster size, suggesting the formation of a primary interaction while retaining structural flexibility. For comparison, the MMP-9–CCR3 complex exhibited a relatively more stable cluster, with a HADDOCK score of -83.1 ± 9.5 and a Z-score of -1.9. The notably low RMSD value indicates a more compact structure with reduced variability, although this may also reflect a smaller number of sampled conformations. These results indicate that MMP-9 establishes more stable interactions with CCR3, whereas MMP-7 engages in more frequent, potentially more flexible interactions. Collectively, these observations support our hypothesis that the MMP-9–CCR3 axis constitutes a critical molecular pathway in the pathogenesis of RSV infection. The 3D visualization further corroborates the presence of a consistent interaction interface in both complexes, reinforcing the biological plausibility of these interactions.

**Figure 2.**
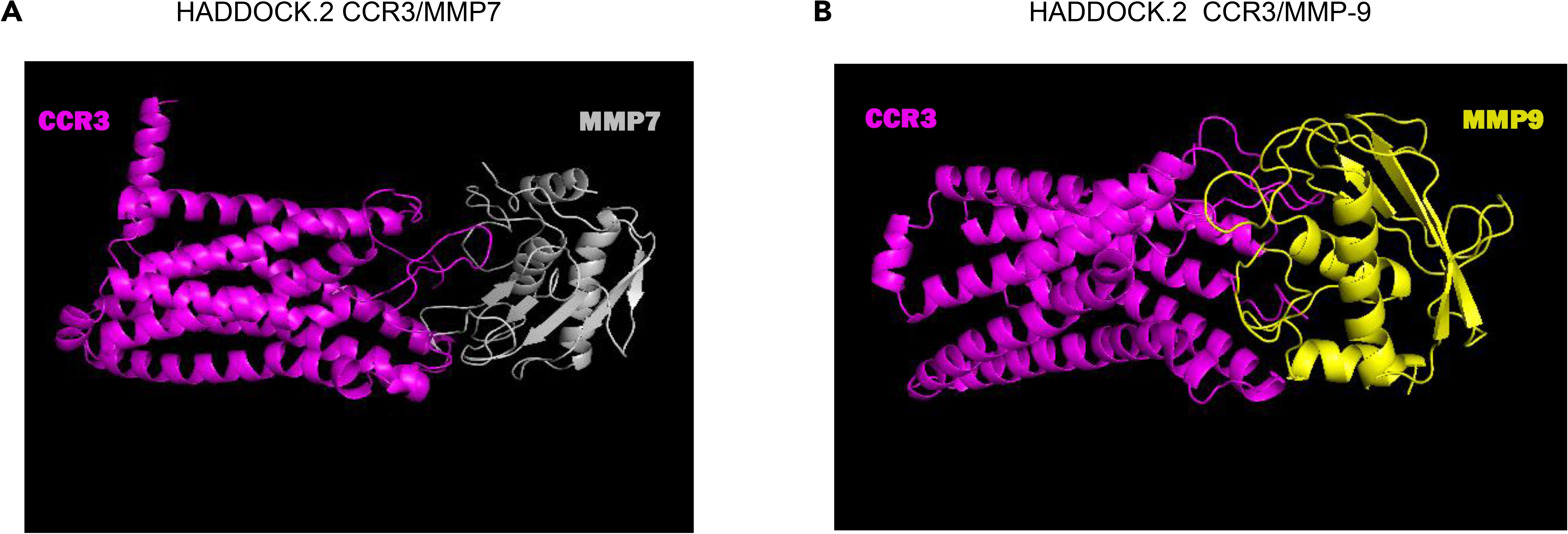
Docking of CCR3 by MMPs via HADDOCK.2. Structural modeling of interactions between matrix metalloproteinases (MMP-9 in yellow, MMP-7 in grey) and the CCR3 receptor (in magenta) generated using HADDOCK2. The figure shows two predicted complexes: A: MMP-9–CCR3 and B: MMP-7–CCR3, illustrating their 3D conformations and potential interaction interfaces.

### Degradation of CCR3 by MMPs

Similar to other G protein-coupled receptors, the N-terminal extracellular region of CCR3 constitutes an exposed domain that is susceptible to the proteolytic cleavage by proteinases^46^. Because the extracellular N terminus of CCR3 is a major determinant for high-affinity binding of eotaxin-1/CCL11, we investigated whether CCR3 is prone to cleavage by metalloproteinases. In a first set of experiments, the effect of MMPs on the cell surface expression of CCR3 was examined in CCR3-transfected Ghost-cells (G.CCR3^+^) and CCR5-transfected Ghost-cells (G.CCR5^+^) (Figure 3A). Electrophoretic separation of proteins followed by Western blotting using an antibody against CCR3 was performed to obtain molecular evidence of CCR3 cleavage by MMPs. A protein band with a relative molecular mass consistent with the expected size of intact CCR3 (∼45 kDa) was detected in cells incubated with medium alone or treated with MMP-1 (Figure 3A), indicating no cleavage^47^. In contrast, Western blot analysis revealed that treatment of CCR3 cells with MMP-7 generated a lower–molecular–weight form of CCR3 (Figure 3A), demonstrating that CCR3 is a novel substrate for MMP-7. MMP-12 was also capable of cleaving CCR3 (Figure 3A). Similar results were obtained using recombinant pro–matrilysin activated with 1 mM APMA (data not shown). In contrast, CCR5, another CC chemokine receptor that binds MIP-1β/CCL4, was not cleaved by MMP-1, MMP-7, or MMP-12 (Figure 3 and data not shown). Increasing concentrations of the MMP-7 catalytic domain (1, 5, 10, 25, and 100 ng/ml) were tested to determine the optimal dose for CCR3 cleavage; the maximal effective concentration was 25 ng/ml (Figure 3C), which was subsequently used in all experiments. The broad-spectrum MMP inhibitor GM6001 effectively preserved CCR3 expression by preventing MMP-7–mediated proteolytic cleavage (Figure 3D)

**Figure 3:**
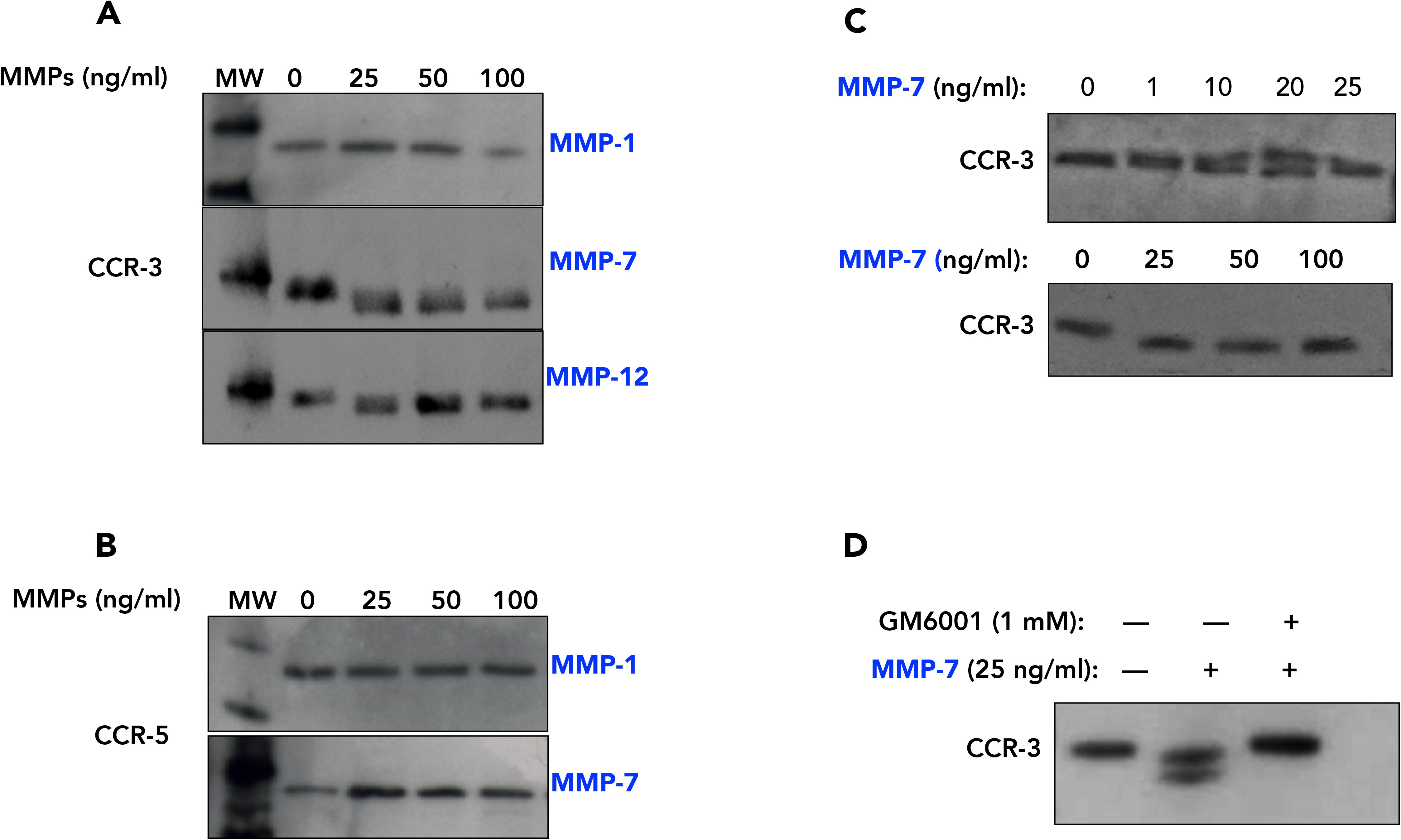
Specificity of MMP cleavage of CCR3. Ghost cells transfected with CCR3 (A) or CCR5 (B) were incubated with or without MMPs (25, 50, 100 ng/ml) at 37°C for 4 h. Molecular weights of the chemokine receptors were determined by Tris-tricine SDS-PAGE and immunoblotting as described in Materials and Methods. A. Unaltered biochemical properties of CCR3 obtained in the presence of medium (alone) or MMP-1was verified by tricine-gel and immunoblotting. CCR3 truncated by MMP-7 and MMP-12 had reduced molecular weight compared to that of intact CCR3. B. CCR5 (anther member of CC chemokine receptor family) is not affected by all MMPs tested. C. The cleavage of CCR3 by MMP-7 was maintained as lower concentrations but optimal at 25 ng/ml of MMP-7. D. The addition of GM6001 (1 mM) together with MMP-7 inhibits the cleavage of CCR3. MW: mass of molecular weight markers

### MMP-7 treatment of CCR3 cells reduce the binding of eotaxin/CCL11 to CCR3

To assess the biological activity of the MMP-7–cleaved form of CCR3, we evaluated the binding of eotaxin-1/CCL11 to CCR3-expressing cells by flow cytometry analysis (Figure 4A–F). G.CCR3 cells were treated with different MMPs, and the binding of biotinylated eotaxin-1/CCL11 was detected using avidin–FITC. The binding assays demonstrated that no significant change in eotaxin-1/CCL11 binding was detected when cells were treated with MMP-1, MMP-2, MMP-9, or MMP-10, suggesting that these MMPs do not cleave CCR3 in a manner that affects its ligand-binding capacity (Figure 4A–D). However, eotaxin-1/CCL11 binding to MMP-7–cleaved CCR3 was significantly reduced, indicating that MMP-7–mediated proteolysis impairs the receptor’s ability to bind its ligand (Figure 4E). A comparable reduction in binding was also observed following MMP-12 treatment (Figure 4F). CCR5-transfected cells were also used in view to assess the susceptibility of this chemokine receptor to MMP-7, MMP-10, and MMP-12 under identical conditions. MMP treatment did not affect MIP-1β/CCL4 binding to CCR5, confirming that the observed effects on eotaxin-1/CCL11 binding were specific to CCR3 (Figure 4G–I). In a complementary experiment, eosinophils were cultured either in medium alone or in the presence of MMP-7 during 4h, and their ability to express CCR3 and bind eotaxin-1/CCL11 was assessed after 24 hours of incubation. Eosinophils maintained in culture medium readily expressed CCR3 and bound eotaxin-1/CCL11, whereas MMP-7–treated eosinophils showed reduced, but not completely abolished, ligand binding (data not shown). These results suggest that MMP-7–mediated proteolysis partially impairs CCR3 function without fully eliminating its synthesis or its ability to engage the eotaxin-1/CCL11 ligand.

**Figure 4:**
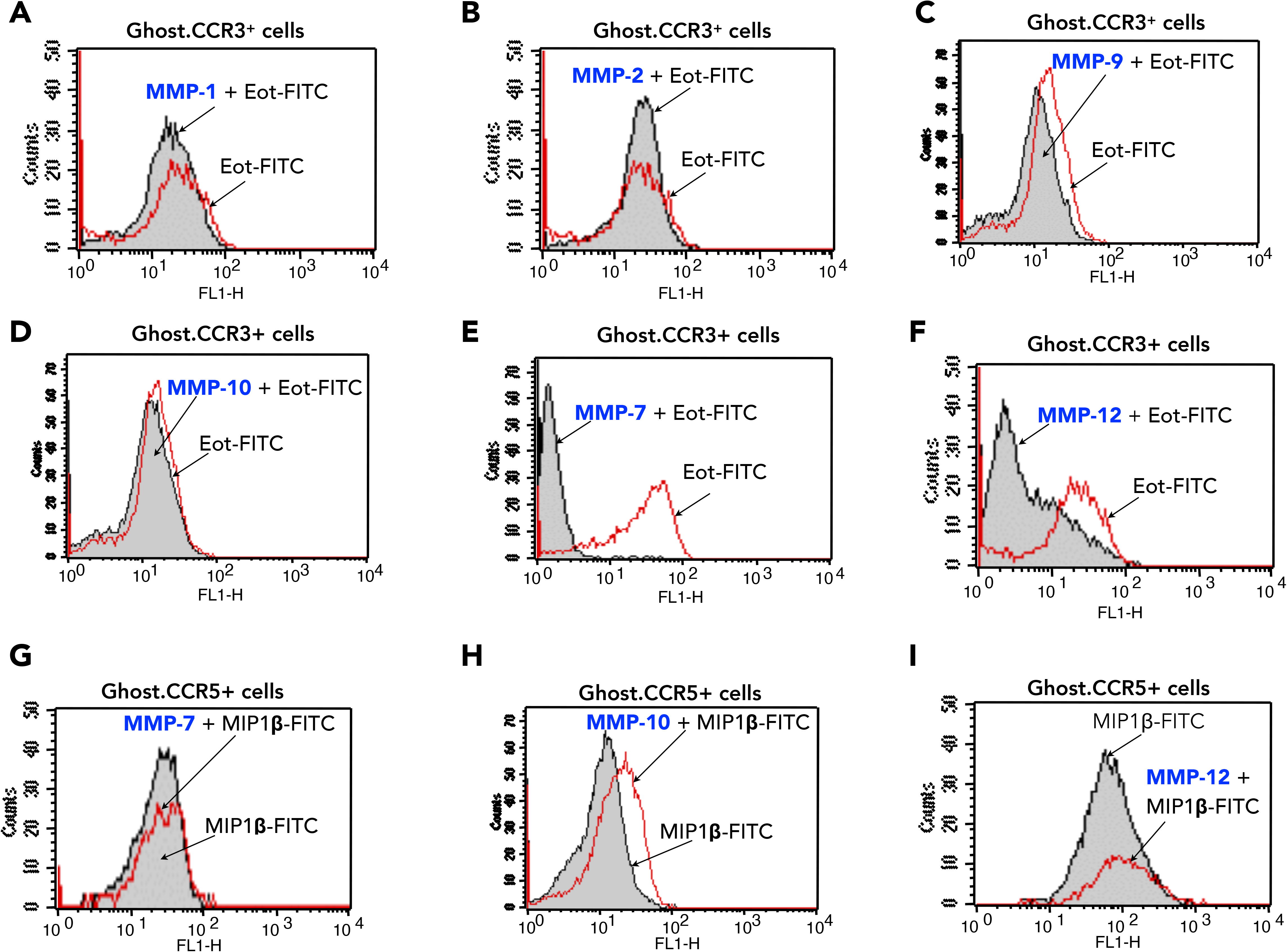
MMP-7 cleaved CCR3 loses affinity for eotaxin-1/CCL11 in CCR3 transfected cells. Proteolytic cleavage by recombinant MMPs was performed using CCR3 or CCR5-Ghost transfected cells. Eotaxin-1/CCL11 or MIP-1β/CCL4 binding to Ghost CCR3^+^ or CCR5^+^ cells in the presence or absence of MMPs were determined by flow cytometry using biotin labeled eotaxin-1/CCL11 or biotin labeled MIP-1β/CCL4 followed by streptavidin-FITC conjugate (R&D Systems). Binding of eotaxin-1/CCL11 to its receptor was unaltered by MMP1, -2, -9 or –10 (A-D), whereas treatment of CCR3-Ghost cells with MMP-7 (E) or MMP12 (F) reduced the binding of eotaxin-1/CCL11 to its receptor. Binding of MIP1β/CCL4 to CCR5 was unaltered by all the MMPs tested (G-I). However, MMP10 treatment of GHOST-CCR5 cells increased slightly the binding of MIP1β/CCL4 to its receptor.

### CCR3 cleaved form lost binding to eotaxin-1/CCL11 in eosinophils

The consequence of CCR3 degradation on the eotaxin-1/CLL11 binding was next examined in eosinophils, which naturally express high levels of CCR3. Cells were exposed to rhMMP-7, and CCR3 cleavage was evaluated by measuring the binding of biotinylated eotaxin-1/CLL11 detected with avidin-FITC (Figure 5A) or by antibody labelling followed by flow cytometry analysis (Figure 5B). MMP-7–treated eosinophils exhibited a marked loss of eotaxin-1/CCL11 binding, indicating that proteolytic cleavage of CCR3 profoundly impairs ligand recognition (Figure 5A). This loss of binding is consistent with the removal of the eotaxin-1/CCL11 binding motif located within the CCR3 N terminus, a region critical for high-affinity ligand interaction. Supporting this interpretation, monoclonal antibodies directed against the CCR3 amino-terminal domain also failed to recognize the cleaved receptor (Figure 5B), ruling out steric interference or competitive binding by MMP-7 as alternative explanations.

**Figure 5:**
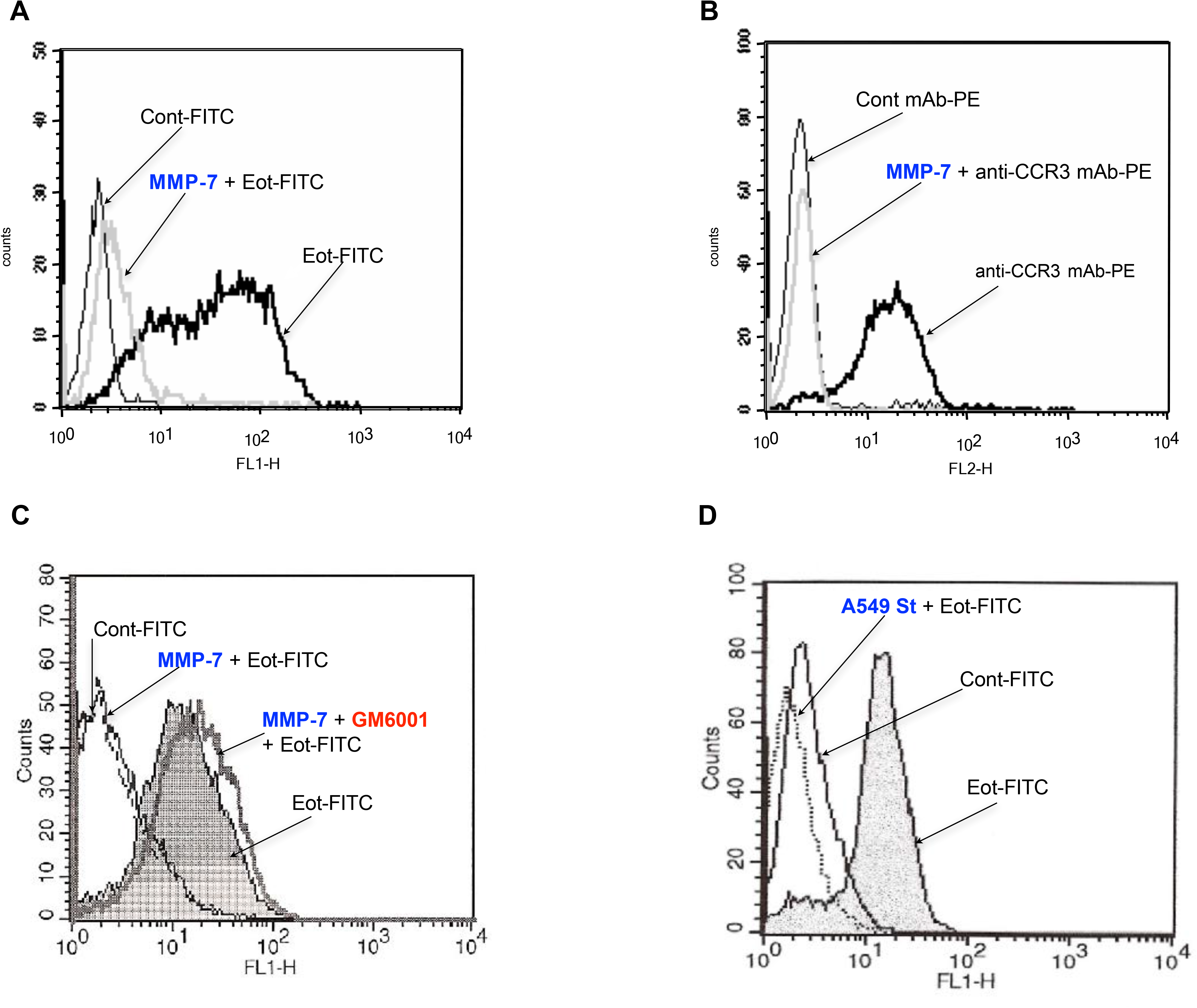
MMP-7 cleaved CCR3 loses affinity for eotaxin-1/CCL11 binding in eosinophils. Eosinophils (4 x 10^6^ cells/ml) were exposed or not to MMP-7 for 4 h at 37 °C prior to incubation with biotinylated eotaxin-1/CCL11 (A) or PE-labelled anti-CCR3 Abs (B). In the case of eotaxin-1/CCL11, cells were further incubated with the avidin-FITC reagent for an additional 30 min at 4°C in the dark. Flow cytometry analysis revealed that MMP-7 pretreatment of eosinophils impaired the binding of both eotaxin-1/CCL11 and anti-CCR3 Abs to CCR3. The activity of MMP-7 was inhibited by GM6001 and allows binding of eotaxin-1/CCL11 to CCR3 (C). Conditioned media derived from cultured A549 cells, used as a source of MMPs, prevented eotaxin-1/CCL11 binding to CCR3 eosinophils (D).

Importantly, inhibition of MMP-7 enzymatic activity with GM6001, a potent and reversible broad-spectrum inhibitor of MMPs, restored eotaxin-1/CCL11 binding as demonstrated by FACS staining (Figure 5C), confirming that the loss of ligand recognition was dependent on MMP-7 catalytic activity. To enhance the physiological relevance of these findings, supernatants from A549 airway epithelial cells, known to express MMP-7, were used in place of recombinant enzyme. The A549 supernatant similarly reduced eotaxin-1/CCL11 binding to CCR3 (Figure 5D), demonstrating that CCR3 cleavage is undoubtedly mediated by the catalytic activity of matrilysin under physiologically relevant conditions.

### Broader impact of MMP-7 on chemokine receptor integrity

To determine whether MMP-7–mediated cleavage extends to other chemokine receptors, we examined the susceptibility of additional receptors to MMP-7–induced proteolysis. G.CCR1, G.CCR5, G.CXCR4, and G.CX3CR1 cells were treated with MMP-7, and receptor cleavage was assessed using ligand-binding assays and antibody labelling, followed by flow cytometric analysis. MMP-7 treatment of G.CCR1 cells resulted in a significant reduction in MIP-1α/CCL3 binding, indicating that MMP-7 impairs CCR1 ligand-binding capacity probably through cleavage of CCR1 (Figure 6A). Similarly, exposure of G.CXCR4 cells to MMP-7 caused a pronounced decrease in SDF-1/CXCL12 binding (Figure 6B). In contrast, treatment of G.CCR5 cells with MMP-7 slightly enhanced MIP-1β/CCL4 binding (Figure 6C), suggesting a possible conformational modification rather than proteolytic inactivation. These findings indicate that MMP-7 selectively cleaves CCR1 and CXCR4, thereby disrupting their ability to bind their respective ligands, whereas its effect on CCR5 is minimal and may involve subtle conformational changes that do not impair ligand recognition. In contrast, MMP-7 treatment had no effect on the binding of an anti-CX3CR1 monoclonal antibody directed against an extracellular loop epitope (Figure 6D), suggesting that CX3CR1 is resistant to MMP-7–mediated proteolysis. Collectively, these results demonstrate that MMP-7 may targets specific chemokine receptors, such as CCR1 and CXCR4, while sparing others like CX3CR1. The cleavage of these receptors and its functional consequences are under investigation.

**Figure 6:**
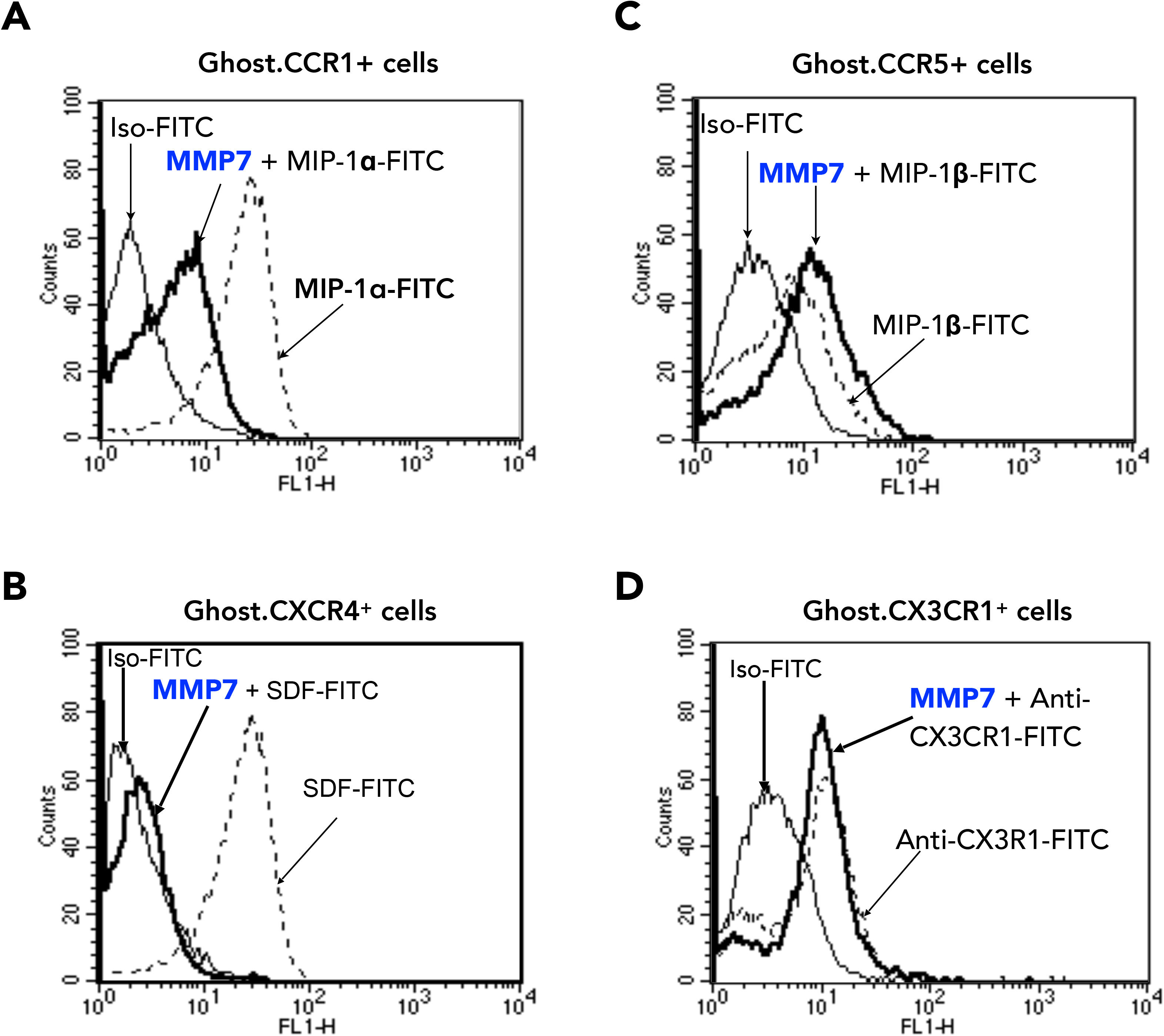
Effect of MMP-7 on CCR1, CCR5, CXCR4 and CX3CR1 chemokine receptors. CCR1, CCR3, CCR5, CXCR4 and CX3CR1 transfected cells (4 x 10^6^ cells/ml) were exposed to MMP-7 for 4 hours at 37°C and cleavage of these chemokine receptors were followed by flow cytometry analysis. The expression of CCR1, CCR3, CCR5 and CXCR4 were assessed with biotinylated ligands (MIP-1 /CCL3, eotaxin-1/CCL11, MIP-1 /CCL4 and SDF/CXCL12) and avidin-FITC. In the case of CX3CR1, cells were stained with anti-CX3CR1 Abs. MMP-7 treatment of CCR1 and CXCR4 transfected cells reduced their binding to the corresponding ligands (MIP-1α/CCL3 and SDF/CXCL12 respectively) (A and B) and has no effect on the binding of MIP-1 /CCL4 to CCR5 (C). In contrast, binding of anti CX3CR1mAb, which recognize an epitope located in the extracellular loop of CX_3_CR1, remained unaffected (D). The binding of eotaxin-1/CCL11 or anti-CCR3-PE antibody to CCR3+ cells was reduced following MM-7 treatment of Ghost-CCR3+ cells (E, F).

### MMP-7–driven CCR3 cleavage impairs CCR3 cell chemotaxis

Binding of chemokines to their corresponding receptors is a critical step in initiating downstream cellular responses such as chemotaxis. To evaluate the functional impact of MMP-7–induced CCR3 cleavage, we compared the ability of eotaxin-1/CCL11 to induce chemotaxis in MMP-7–treated and untreated cells using Boyden chamber assays. Pretreatment of eosinophils and G.CCR3 cells with MMP-7 or MMP-12 for 4 hours at 37 °C markedly reduced their chemotactic response to eotaxin-1/CCL11 (Figure 7A and 7B). In contrast, treatment with MMP-1, MMP-2, MMP-3, MMP-9, or MMP-13 had no detectable effect on migration (Figure 7A and data not shown). Similarly, MMP-7 treatment inhibited chemotaxis in G.CCR1 and G.CXCR4 cells, whereas no significant changes were observed in G.CCR5 or G.CX3CR1 cells under the same conditions (Figure 7B).

**Figure 7:**
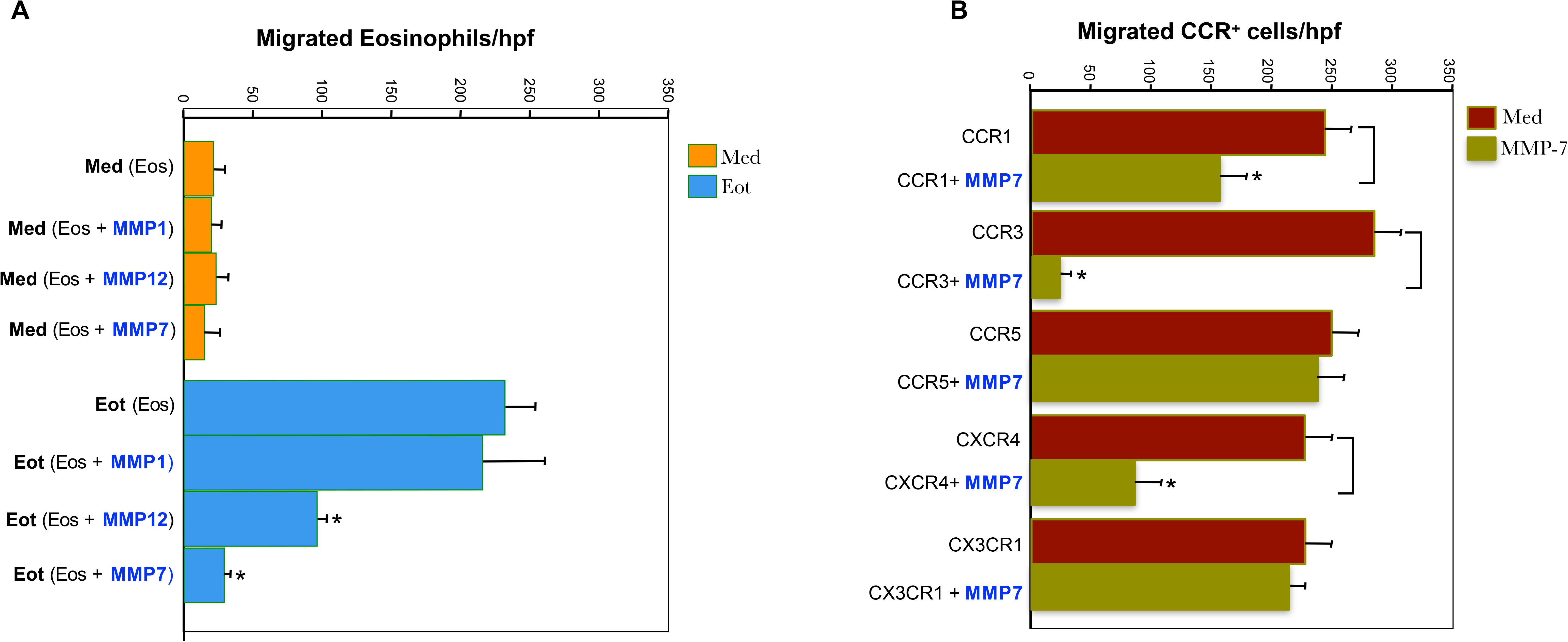
Migration assay of cells treated by MMP-7. Dose-response curves of cellular migration induced by eotaxin-1/CCL11 (Eot) in filter assays and inhibition of chemotactic activity of eotaxin-1/CCL11 by MMP-7 and -12. (A) Eosinophils (Eos) were preincubated with medium alone, MMP-1, MMP-7 or MMP-12 during 4 h at 37°C and a chemotaxis assay was performed with eotaxin-1/CCL11 at different concentrations in a Boyden chamber. (B) CCRs transfected Ghost cells were preincubated with medium alone or MMP-7 during 4h at 37°C and a chemotaxis assay was performed with the corresponding ligands (MIP-1α/CCL3, Eotaxin-1/CCL11, MIP-1 /CCL4, SDF/CXCL12 and Fractalkine) in a Boyden chamber. Data are presented as the number of cells per high power field (magnification, X400). The results shown are representative experiments and are presented as the mean ± SD of 5 fields per well.

Eotaxin-1/CCL11 is known to activate eosinophils through CCR3^37^. To establish whether the activation of ERK2 by eotaxin-1/CCL11 was affected by proteases, eosinophils were preincubated with MMP-7 (25 ng/mL) for 4 hours before stimulation with eotaxin-1/CCL11 (10 ng/ml) for 1 minute. As assessed by Western blotting using antibodies against dual-phosphorylated ERK2, cells pretreated with MMP-7 (25 ng/ml) for 4 hours prior to eotaxin-1/CCL11 stimulation completely abrogated activation of extracellular signal-regulated kinase 2 (ERK2), a key mitogen-activated protein (MAP) kinase in the CCR3 signaling pathway (Figure S1). Stimulation of eosinophils with MMP-7 alone (25 or 100 ng/mL) did not elicit phosphorylation of ERK (data not shown). Taken together, these results indicate that MMP-7 treatment not only impairs chemotaxis but also disrupts CCR3-mediated intracellular signaling pathways that are essential for directed cell migration.

### MMP-7-processed CCR3 limits RSV infection of airway epithelial cells

CCR3 has been reported by our group and suggested by others to serve as a receptor for RSV ^5,48,49^. To investigate whether MMP-7 modulates RSV infection, plaque assays were performed to assess viral infectivity in airway epithelial cells. The effects of various MMPs were compared for their ability to reduce RSV infection in A549 cells, which express CCR3^47^. As shown in Figure 8, treatment of A549 cell monolayers with increasing concentrations of recombinant MMP-7 led to a significant reduction in RSV infectivity, indicating that MMP-7–mediated cleavage of CCR3 can impair RSV entry into host cells. At the highest concentration tested (500 ng/ml), MMP-7 treatment reduced RSV infection to approximately 6% of control levels, where control cells were incubated with virus in medium alone (Figure 8A). Similar inhibitory effects were observed in Hep-2 cells (data not shown). Treatment with heparin, previously reported to inhibit RSV infection, produced comparable levels of viral inhibition (Figure 8A) ^41^. The inhibitory effect of MMP-7 was dose-dependent, with lower concentrations (1, 5, and 10 ng/ml) showing progressively less inhibition, while 50 ng/ml MMP-7 markedly reduced infection (Figure 8B). Although MMP-12 also cleaves CCR3 and impairs eotaxin-1/CCL11 binding and chemotaxis, it had no effect on RSV infection of airway epithelial cells (Figure 8A, B). This may reflect differences in the CCR3 cleavage sites targeted by MMP-7 versus MMP-12. Exposure of A549 cells to MMP-1 did not impair RSV infectivity (Figure 8A, B). Furthermore, MMP-7 treatment inhibited RSV infection in CCR3-transfected cells and in Hep-2 cells, whereas none of the other MMPs tested showed inhibitory activity at comparable concentrations (data not shown). Collectively, these findings indicate that MMP-7 specifically modulates RSV infection by processing CCR3, thereby limiting viral entry into airway epithelial cells.

**Figure 8.**
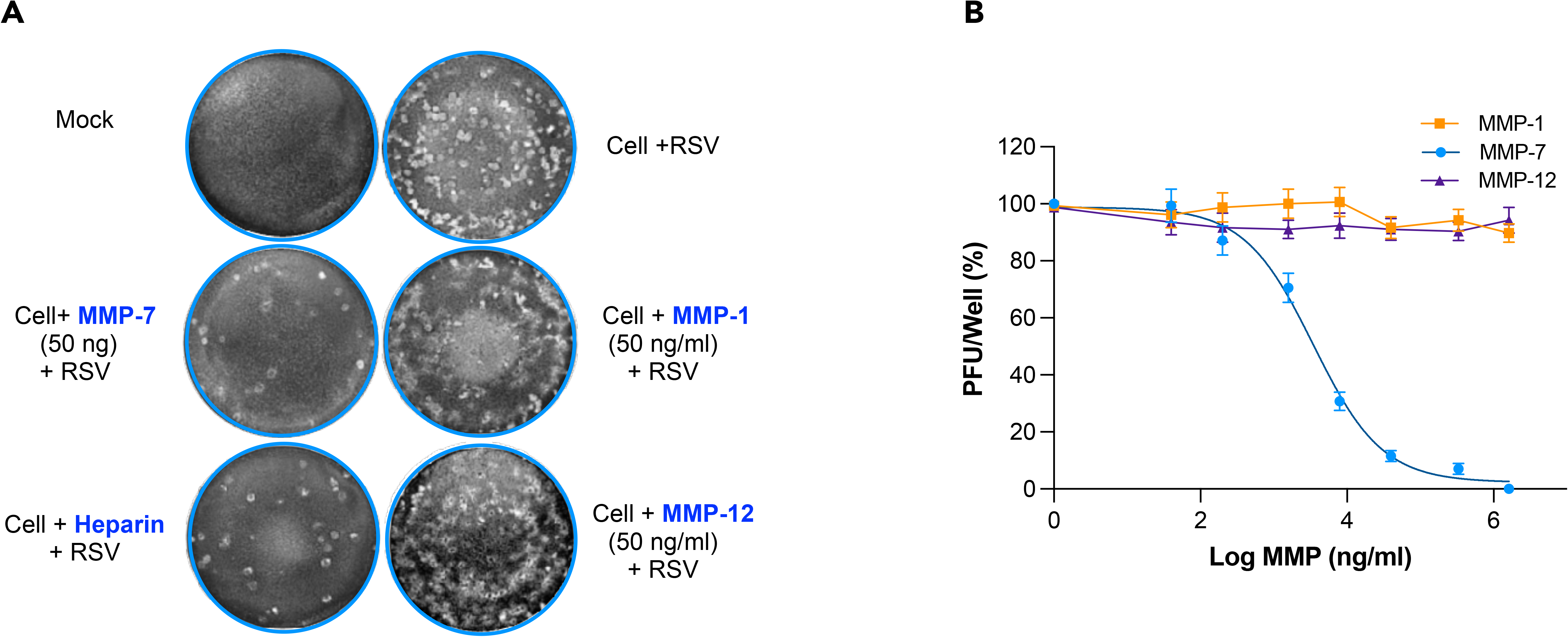
MMP-7 inhibits infection of human epithelial cells with RSV. MMP-1, -7, -12 were assessed for their ability to block CCR3 dependent RSV infection, which revealed no inhibition by MMP-1 and -12 in contrast to MMP-7. A549 cells were exposed to MMP-1, -7, and -12 at different concentration for 4 hours at 37°C. Thereafter, the cells were rinsed three times with culture medium and infected for 1h with A2 strain of RSV. After 3 to 4 days incubation, plaques were enumerated to determine the degree of viral infectivity. Focus forming units per well were then counted using light microscopy. (A) Plaque morphology of RSV A2 strain after 3 to 4 days incubation with A549 cells pretreated with MMP-1, -7, -12 (50 ng/ml) or heparin (5 µg/ml). (B) Normalization, GraphPad. All error bars indicate mean ± SD of three independent experiments performed. *P<0.05 versus MMPs-treated cells; unpaired t-test.

### Docking Analysis of RSV G protein with MMP-7 via ColabFold2: Structural Stability and Interaction Insights

To assess the potential interaction between MMP-7 and the RSV G protein, structural docking analyses were performed (Figure S2A). One alignment showed higher predicted stability than an alternative MMP-7–G configuration, with an approximately 50% increase in predictive confidence. Structural confidence was evaluated using pLDDT color mapping, which indicated that the majority of protein regions adopt well-defined conformations (blue and yellow), corresponding to a mean confidence score of 80.38%. Terminal segments and loop regions exhibited lower confidence scores, represented in red, reflecting intrinsic flexibility or disorder and accounting for approximately 19.62% of the overall structure. Additionally, the sequence coverage profile showed a high degree of homogeneity and conservation, further reinforcing the structural reliability of the predicted complex.

A comparative analysis was performed for the RSV G protein in complex with MMP-9 (Figure S1B). This interaction exhibited moderate structural stability, as indicated by pLDDT scores and sequence alignment coverage. Well-defined regions, shown in blue with scores above 80.77%, reflect areas of high structural confidence, whereas yellow, orange, and red regions—accounting for approximately 19.23% of the structure—indicate increased flexibility or disorder. The sequence coverage graph further supports this interpretation, showing conserved domains covering roughly 70% of the sequence, which enhances the reliability of the predicted structure. Overall, these results indicate that both RSV-G–MMP-7 and RSV-G–MMP-9 complexes adopt predominantly stable conformations, while retaining flexible domains that may facilitate dynamic interactions, including proteolytic processing.

### Molecular Docking Analysis of RSV G protein with MMP-7 and MMP-9 via HADDOCK

Initial in silico screening suggested a potential role for MMPs in modulating the RSV G protein. To validate these predictions, semi-flexible docking was performed using HADDOCK 2.4. The main clusters obtained for the MMP-9–G protein complex exhibited average binding energies ranging from –111.9 ± 8.5 kcal/mol to –99.1 ± 5.6 kcal/mol. Cluster 1, representing the most populated and favorable cluster, showed the lowest interface energy, indicative of a stable and potentially functional interaction. Analysis of the interaction interfaces supports the likelihood of direct contact between MMP-9 and the RSV G protein, with high model reliability (Figure S3).

Docking analysis of MMP-7 with the RSV G protein revealed an energetically favorable interaction. Cluster 15, although modest in size, exhibited an average binding energy of –119.5 ± 5.1 kcal/mol and an interface energy of 0.4 ± 0.2 kcal/mol, indicative of a strong and stable interaction. Cluster 2 also displayed favorable binding with a conserved interaction interface, supporting the hypothesis of a specific MMP-7–G protein interaction (Figure S3). Comparison with MMP-9 indicates that MMP-7 exhibits slightly higher binding affinity, whereas MMP-9 forms a more stable and populated conformation, making it a credible target as well. In both cases, the interaction interfaces were validated, confirming the biological plausibility of these complexes. Overall, these results support the initial predictive docking models and suggest that RSV G protein could be cleaved or modulated by MMP-7 and MMP-9, potentially influencing RSV infectivity.

### MMP-7 Cleavage of RSV-G protein reduces viral infectivity in airway epithelial cells

These experiments highlight the role of matrix metalloproteinases (MMPs), particularly MMP-7, in modulating RSV infectivity through interactions with the viral RSV G protein. RSV G, an essential attachment glycoprotein, mediates viral entry by binding to host cell receptors. Proteolytic assays demonstrated that both MMP-7 and MMP-9 cleave RSV G, as evidenced by reduced band intensity and altered molecular weight, consistent with enzymatic degradation (Figure 9A). This cleavage likely compromises RSV G structural integrity, impairing its capacity to mediate viral attachment and infection. Furthermore, dose-dependent reductions in viral plaque formation with increasing MMP-7 concentrations reinforced its inhibitory effect on RSV infectivity (Figure 9B–D). Quantitative analysis confirmed that MMP-7 significantly reduces RSV infectivity. Together, these findings indicate that MMP-7 interacts with RSV G and is associated with interference in viral processes relevant to infection.

**Figure 9:**
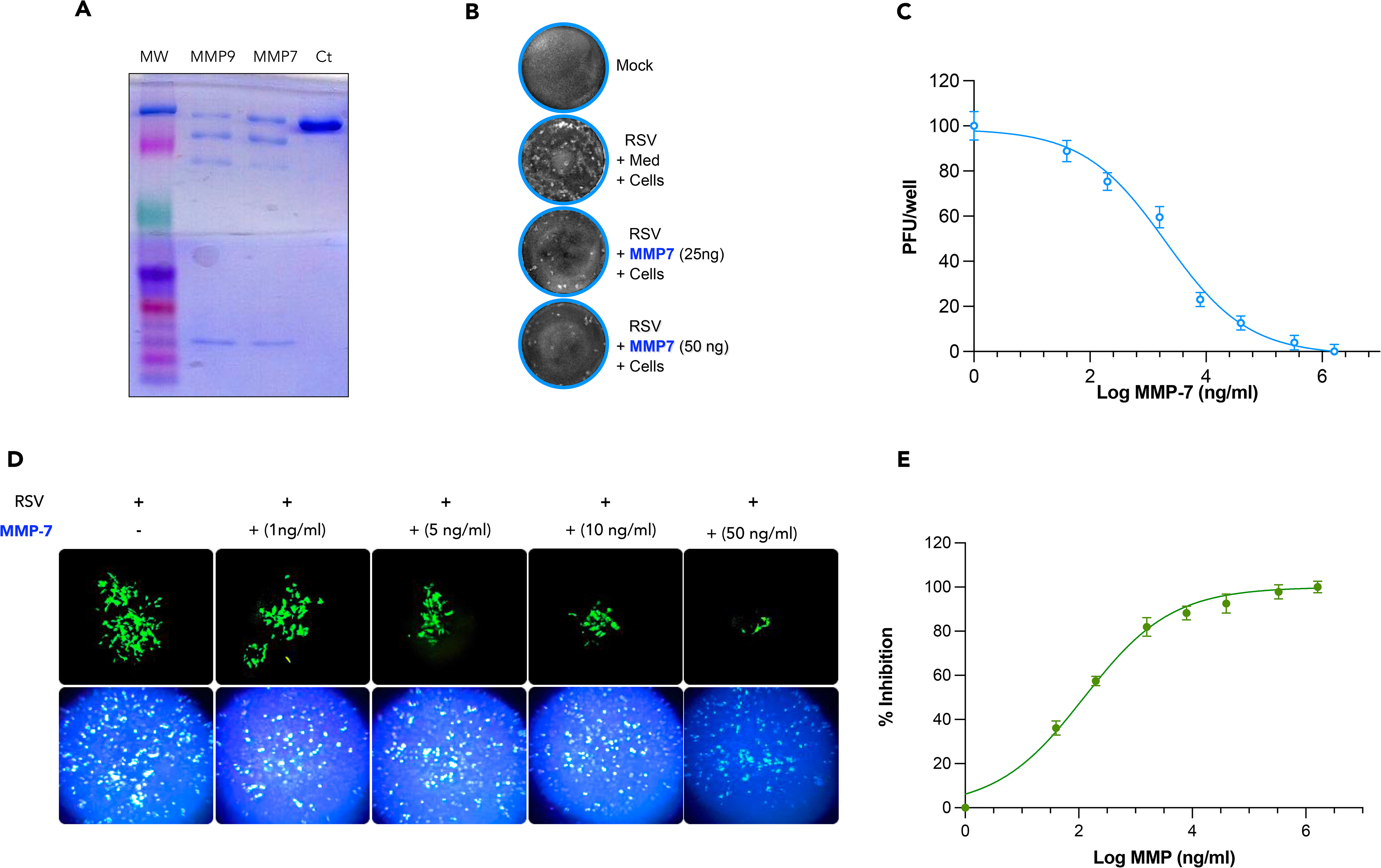
MMP-7 Cleaves RSV G protein and Reduces RSV Infectivity. (A) Analysis of RSV G protein, with coomassie-stained gel showing a band corresponding to Gp90. Incubation with MMP-7 or MMP-9 shows degradation of the Gp90 protein, with visible changes in band intensity and molecular weight patterns. (B) Plaque assay results demonstrating reduced plaque formation when RSV was treated with increasing concentrations of MMP-7, 25 ng and 50 ng, compared to the medium (Med) treated RSV and Mock treated cells. (C) Quantitative bar graph showing % inhibition of RSV infectivity by various concentrations of MMP-7. (D) Fluorescent images show strong RSV-GFP spread without treatment. Higher MMP-7 reduced green signal, indicating less infection. Blue DAPI staining shows equal cell numbers. The bottom graph shows the corresponding percent inhibition, which increases with rising MMP-7 concentration.

### Increased Susceptibility of MMP-7^□/□^ Mice to RSV Infection

To investigate the functional consequences of MMP-7 deficiency during RSV infection, viral replication and pulmonary pathology were assessed in MMP-7□/□mice. Quantitative analysis of viral load revealed that MMP-7^□/□^ mice harbored significantly higher levels of RSV in lung tissue compared with wild-type (WT) controls (Figure 10A), indicating that MMP-7 contributes to viral clearance or limits replication within the lungs. Consistent with these findings, ELISA measurements showed elevated RSV antigen concentrations in MMP-7^□/□^ mice relative to WT animals. Histological analysis further highlighted the pathological consequences of MMP-7 deficiency (Figure 10B). H&E staining revealed disrupted lung architecture and increased infiltration of inflammatory cells in MMP-7^□/□^ mice compared with WT controls, suggesting that MMP-7 may modulate immune cell recruitment or activity during viral infection. PAS staining demonstrated substantial mucus accumulation in MMP-7^□/□^ lungs, indicative of exacerbated mucus hypersecretion. Quantitative assessment using the mucus index confirmed a statistically significant increase in MMP-7^□/□^ mice (*p < 0.05) (Figure 10C). These results collectively highlight the pivotal role of MMP-7 in host defense during RSV infection, likely through both immunomodulatory and direct antiviral mechanisms, and underscore its importance in modulating lung inflammation and mucus production.

**Figure 10.**
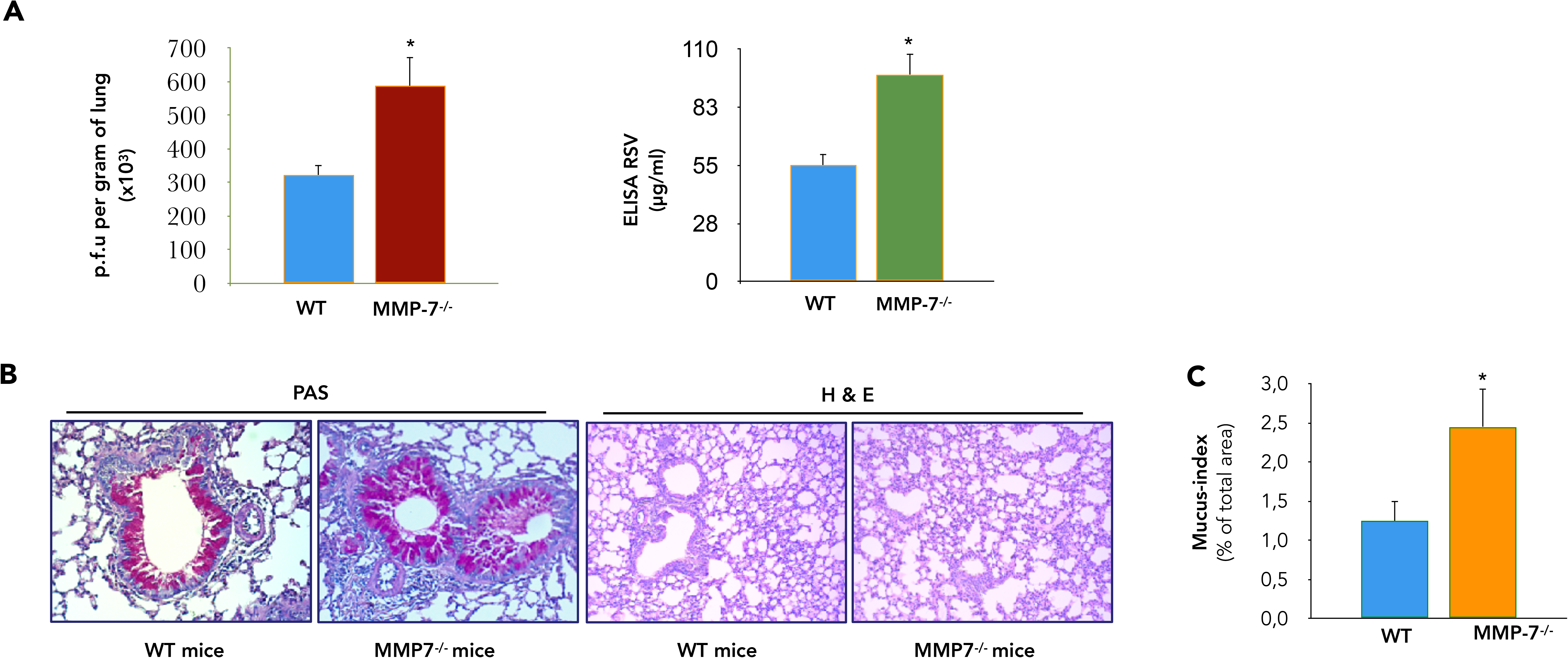
MMP-7 deficiency increases lung RSV-induced inflammation and mucus production. (A) Effect of MMP-7 deficiency on lung viral titer 5 d after infection. Animals were infected, and 5 days post-infection they were killed. Bar graphs show quantitative plaque assays and ELISA of RSV in lung homogenates. P<0.001, n=7, and n=8 per group for experiments in RSV plaque assay and Elisa respectively. (B) Lungs from RSV-infected wild type and MMP-7^-/-^ mice were harvested 5 dpi and then sectioned. Paraffin-embedded lung tissues obtained from RSV-infected wild type and MMP-7^-/-^ mice were stained by Hematoxyline and Eosine (H&E) and periodic acid-Schiff (PAS). Lungs show increased inflammation in MMP-7 deficient mice and mucus production as evidenced by significant fewer PAS-positive cells. (C) Bar graph shows the percentage of PAS positive cells as the ratio of airway epithelial cells positive for mucus production over the total number of airway epithelial cells counted per mouse lung section, multiplied by 100.

## DISCUSSION

Emerging evidence indicates that specific MMPs actively shape the inflammatory milieu by proteolytically processing non-matrix substrates, including cytokines, chemokines, and antimicrobial peptides, thereby modulating their biological activity. Herein, we identify MMP-7—a protease constitutively expressed by airway epithelial cells^43^ —as a regulator of airway inflammation and anti-viral defense. MMP-7 limits eosinophil recruitment by cleaving CCR3, the principal receptor for eotaxin-1 (CCL11), thereby directly dampening chemokine-driven type 2 inflammation. MMP-7 limits eosinophil recruitment by cleaving CCR3, the principal receptor for eotaxin-1 (CCL11), thereby directly dampening chemokine-driven type 2 inflammation. Cleavage of CCR3 also renders airway epithelial cells less permissive to RSV infection, linking MMP-7 activity to both immune cell trafficking and viral susceptibility. Beyond host substrates, MMP-7 cleaves the RSV G attachment glycoprotein, further reducing infection of host cells. Thus, coordinated proteolytic targeting of CCR3 and a viral surface protein may represent a dual mechanism by which MMP-7 simultaneously dampens type 2 inflammation while restricting viral entry.

Eotaxin-1/CCL11, a potent eosinophil chemoattractant, is rapidly induced in epithelial cells upon injury and infection^50^. Through the selective expression of its specific receptor, CCR3, human eotaxin-1/CCL11 is a potent chemoattractant for eosinophils^51,52^, basophils^53^, and Th_2_ lymphocytes^54^, all of which are found in tissues undergoing allergic reactions. Consequently, an agent that could disrupt eotaxin-1/CCL11’s interaction with CCR3 is likely to effectively reduce the number of cell migration to the lung and subsequent airway hyperresponsiveness. This study presents investigations into the proteolytic processing of CCR3 by MMP. Both MMP-7 and MMP-12 efficiently cleave CCR3, rendering it an inactive receptor by compromising its ability to bind eotaxin-1/CCL11. Truncated CCR3 displays reduced affinity to eotaxin-1/CCL11, fails to induce ERK1/2 phosphorylation upon eotaxin-1/CCL11 stimulation (Figure S1) and consequently diminishes the intrinsic migratory capacity of CCR3^+^ cells towards an eotaxin-1/CCL11 gradient *in vitro*.

Corroborating these findings, immune-ablation of eotaxin-1/CCL11 models of lung injury substantially reduces tissue damage, presumably by decreasing eosinophil influx^55^. Similarly, in various injury and inflammation models, eosinophil influx and transepithelial migration are markedly reduced in mice lacking CCR3 or treated with anti-CCR3 antibodies^56^. In parallel, other MMPs-active chemokines, including MMP-2, have been shown to cleave MCP-3. Proteolytically cleaved MCP-3 binds to CCR-1, -2, and -3, but no longer induces calcium fluxes or promotes chemotaxis; thereby acting as a broad chemokine antagonist that dampens inflammation and limits macrophages recruitment^57^. Although the *in vivo* relevance of MMP-7-mediated CCR3 processing remains to be established, coordinated proteolytic targeting of both eotaxin-1/CCL11 and its receptor may represent a critical step for dampening cell migration and restraining the accumulation of CCR3^+^ cells at sites of inflammation

MMPs expression is typically limited to tissue remodelling associated with both normal and abnormal biological processes, such as development, inflammation, tumor growth, and repair. MMP-7, unlike many MMPs, is notably expressed by non-injured, noninflamed mucosal epithelium across most adult tissues^43^. This sustained expression in healthy epithelium suggests its involvement in fundamental homeostatic mechanisms, particularly in providing defence against microorganisms. Indeed, in mice, MMP-7 activates intestinal pro--defensins^58,59^. Consequently, MMP-7-deficient mice, lacking mature active ∝-defensins, exhibit an impaired ability to battle enteric pathogens, including pathogenic Escherichia coli and Salmonella typhimurium^58,60^. Our data further demonstrate that MMP-7 cleaves CCR3 and inhibits RSV infection in airway epithelial cells (Figure 8). Although MMPs have not yet been definitively shown to be involved in defence mechanisms against viral infections of the airways, MMP-7, which has been detected in the airway epithelium^60,61,62^, is a strong candidate for this function. It is thus possible that these effects represent an additional pathway through which MMP-7 contribute to innate immune responses. This hypothesis is consistent with findings in mice where widespread production of MMP-7 in the epithelium is maintained by continuous, low-level bacterial exposure^60^. Supporting this concept, MMP-7 is present at almost undetectable levels in mice, but its production is induced in ex-germ-free mice that are colonised with a single species of commensal bacteria^60^. Our data show that MMP-7 deficiency exacerbates RSV-induced lung pathology, resulting in an increased viral burden, intensified inflammatory responses, and excessive mucus production. These pathological features observed in MMP-7^□/□^ mice underscore the enzyme’s crucial role in maintaining respiratory homeostasis during viral infections. By modulating both viral clearance mechanisms and host immune responses, MMP-7 appears to function as a key regulatory factor in limiting RSV-mediated pulmonary damage. Collectively, these findings support the hypothesis that MMP-7 is not only a central modulator of lung immune dynamics but also a promising therapeutic target for the management of severe RSV-induced respiratory disease.

This evidence prompted our investigation into whether MMP-7 could also cleave the attachment (G) protein of RSV. Our hypothesis was founded on a prior study showing RSV G protein cleavage during virus production in Vero cells^63^. This cleavage markedly reduced infectivity in primary human airway epithelial (HAE) cells, the natural targets of RSV infection. The study identifies cathepsin L as the protease responsible, likely occurs during endocytic recycling^63^. Thus, RSV-G protein cleavage impairs viral attachment and reduces the infectivity of Vero cell-derived virions compared to those bearing intact RSV-G protein^63^. The results presented herein elucidates the potential role of MMP-7 in attenuating viral infectivity through the proteolytic cleavage of viral surface glycoproteins, specifically RSV-G protein. Evidence is presented demonstrating that MMP-7’s capacity to bind and proteolytically process viral proteins critical for host cell entry. This proteolytical event resulted in a conformational disruption of the protein and concomitant impairment of its interaction with cellular receptors, as evidenced by a significant reduction in viral infectivity observed in plaque assays and viral quantification tests. Collectively, these findings substantiate the hypothesis that MMP-7 functions as an endogenous antiviral protease within the respiratory mucosa, capable of restricting viral invasion through dismantling critical RSV structural components.

Collectively, our findings show that MMP-7 targets and modifies proteins central to both viral entry and inflammatory signalling, supporting the view that its enzymatic repertoire extends beyond extracellular matrix remodelling to a broader spectrum of biologically relevant substrates. In the case of RSV, cleavage of the G attachment glycoprotein and of CCR3 converges on a single outcome: reduced epithelial infection together with dampened type 2 inflammation. This activity does not appear to be restricted to RSV. In preliminary experiments not presented here, MMP-7 similarly cleaved the SARS-CoV-2 spike protein; how this processing affects viral infectivity and immune evasion is currently under investigation (I.K. et al., unpublished observations). A systematic survey of MMP-7 substrates, and of other proteases such as neutrophil serine proteases, across respiratory viruses was beyond the scope of the present work. Nonetheless, its capacity to cleave both structural viral proteins and host regulatory receptors makes MMP-7 a compelling candidate for therapeutic strategies aimed at limiting pathogen entry while restraining the accompanying inflammatory response.

In conclusion, our findings demonstrate that MMP-7 cleaves CCR3, which may reduce eosinophils migration to the airways and inhibit RSV infection. Moreover, the observed interactions between MMP-7 and viral proteins, including RSV-G protein and the SARS-CoV-2 spike RBD, may represent an important nexus linking innate immune regulation with host defense against respiratory pathogens. Collectively, these data support a role for MMP-7 in modulating immune cell trafficking and antiviral responses.

In conclusion, our findings identify MMP-7 as a protease acting at the interface of chemokine signalling and viral entry in the airway. MMP-7 cleaves CCR3, reducing eosinophil migration to the airways and rendering epithelial cells less permissive to RSV, and it cleaves the RSV G attachment glycoprotein, further limiting infection. *In vivo*, MMP-7 deficiency aggravates RSV-induced lung pathology. Together, these data support a role for MMP-7 in coordinating immune cell trafficking and antiviral defence and identify it as a candidate target in severe RSV disease.

## LIMITATION OF THE STUDY

Our study has limitations. First, our experimental models do not fully capture the complexity of a natural viral infection. We introduced MMP-7 in isolation or at controlled levels, but during an actual infection, MMP-7 is expressed together with many other proteases that could enhance, modify, or counter its effects. In addition, viral proteins may rapidly activate Interferon-Stimulated Genes (ISGs) before MMP-7–mediated repression takes place. The timing and level of MMP-7 expression can also differ across cell types and host conditions during the viral replication cycle. Therefore, studies using live virus and MMP-7 KO animals, combined with single-cell transcriptomics, are needed to measure MMP-7 activity more precisely and define its role in host resistance and transcriptional regulation.

## ACKNOWLEDGMENTS

The authors thank Bouland and Pr M. D. Diebold (Laboratoire Central d’Anatomie et de Cytologie Pathologiques, CHU de Reims, France) for facilitating sample processing and preparation of lung sections. Special thanks are addressed to all volunteers who, by their consent, allowed this study to be performed.

## AUTHOR CONTRIBUTIONS

B.L. conceptualized the study. A.S.G. and B.L. designed the experiments, directed the research, obtained funding and analyzed the data. Investigation: B.L., I.K., H.A.B., and V.W., performed the experiments. R.L., R.M., and M.K., analyzed the data and participated to critical reading of the manuscript. I. K., A.S.G. and B.L., wrote the manuscript.

## SUPPLEMENTAL INFORMATION

**Figure S1. Effect of MMP-7 on ERK2 activation in eotaxin-stimulated eosinophils.** Eosinophils were preincubated with medium or 1, 10, 25, 50 or 100 ng/ml of MMP-7 for 30 minutes and stimulated with medium or eotaxin (100 ng/ml) for 1 minute. Cytosolic extracts underwent Western blotting with the anti-phospho-ERK1/ERK2 antibody. MMP-7 blocked ERK1/2 phosphorylation in a dose-dependent manner (n =3).

**Figure S2. AI-Predicted docking of G protein with MMPs using ColabFold2** (A) Structural predictions of the MMP-9–G complex generated using Colabfold2, showing moderate confidence with a mean pLDDT score of ∼52.06 and extensive low-confidence regions (colored in red), indicating high flexibility. Predicted aligned error plots reveal considerable uncertainty across chain interfaces. (B) In contrast, structural predictions for the MMP-7–G complex show a significantly higher average pLDDT score (∼80.38), with only 19.6% of residues falling into low-confidence regions, and sequence coverage for MMP-7 indicates alignment with ∼70% of conserved domains, supporting a more stable interaction.

**Figure S3. Docking of Gprotein by MMPs via HADDOCK.2** Molecular modeling of the interactions between MMPs and the viral G protein. (A) Binding interaction between the G protein (depicted in orange) and MMP-9 (shown in cyan). (B) Association between matrix metalloproteinase MMP-7 (colored in orange) and G viral protein (represented in green).

