## Supplementary material for "Matrilysin (MMP-7) Restricts RSV Infection and Airway Inflammation through Proteolysis of CCR3 and RSV-G": Suppl Figures

### Supplemental information

**Figure S1. Effect of MMP-7 on ERK2 activation in eotaxin-stimulated eosinophils.**
Eosinophils were preincubated with medium or 1, 10, 25, 50 or 100 ng/ml of MMP-7 for 30 minutes and stimulated with medium or eotaxin (100 ng/ml) for 1 minute. Cytosolic extracts underwent Western blotting with the anti-phospho-ERK1/ERK2 antibody. MMP-7 blocked ERK1/2 phosphorylation in a dose-dependent manner (n =3).

**Figure S2. AI-Predicted docking of G protein with MMPs using ColabFold2**
(A) Structural predictions of the MMP-9–G complex generated using Colabfold2, showing moderate confidence with a mean pLDDT score of ~52.06 and extensive low-confidence regions (colored in red), indicating high flexibility. Predicted aligned error plots reveal considerable uncertainty across chain interfaces. (B) In contrast, structural predictions for the MMP-7–G complex show a significantly higher average pLDDT score (~80.38), with only 19.6% of residues falling into low-confidence regions, and sequence coverage for MMP-7 indicates alignment with ~70% of conserved domains, supporting a more stable interaction.

**Figure S3. Docking of Gprotein by MMPs via HADDOCK.2**
Molecular modeling of the interactions between MMPs and the viral G protein. (A) Binding interaction between the G protein (depicted in orange) and MMP-9 (shown in cyan). (B) Association between matrix metalloproteinase MMP-7 (colored in orange) and G viral protein (represented in green).

**Fig. S1** **Supplemental information**

A

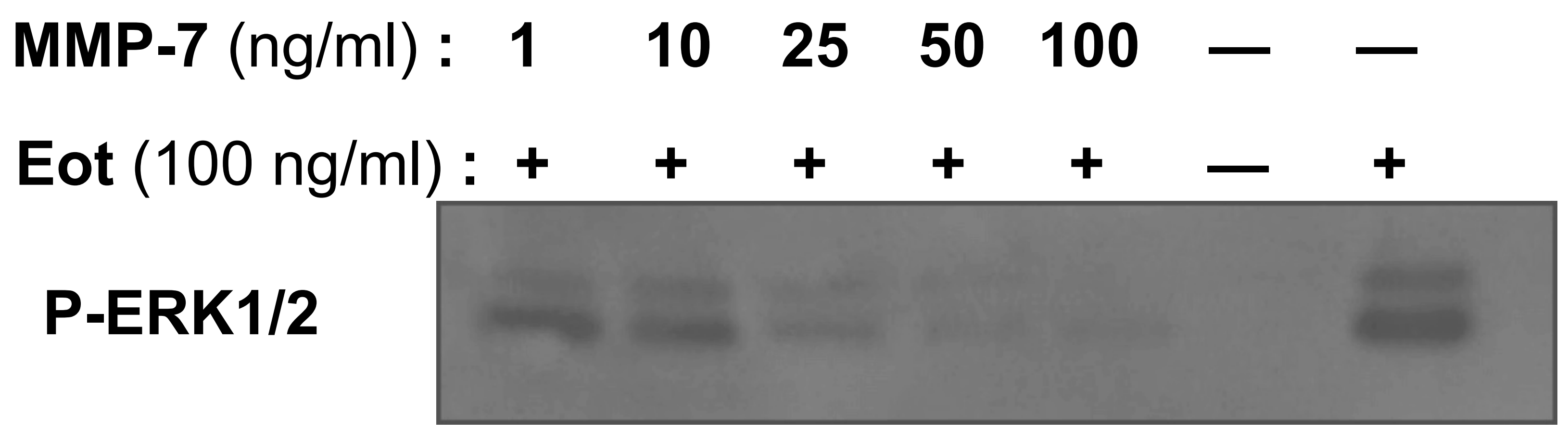

### Supplemental information

**Fig. S2**

A

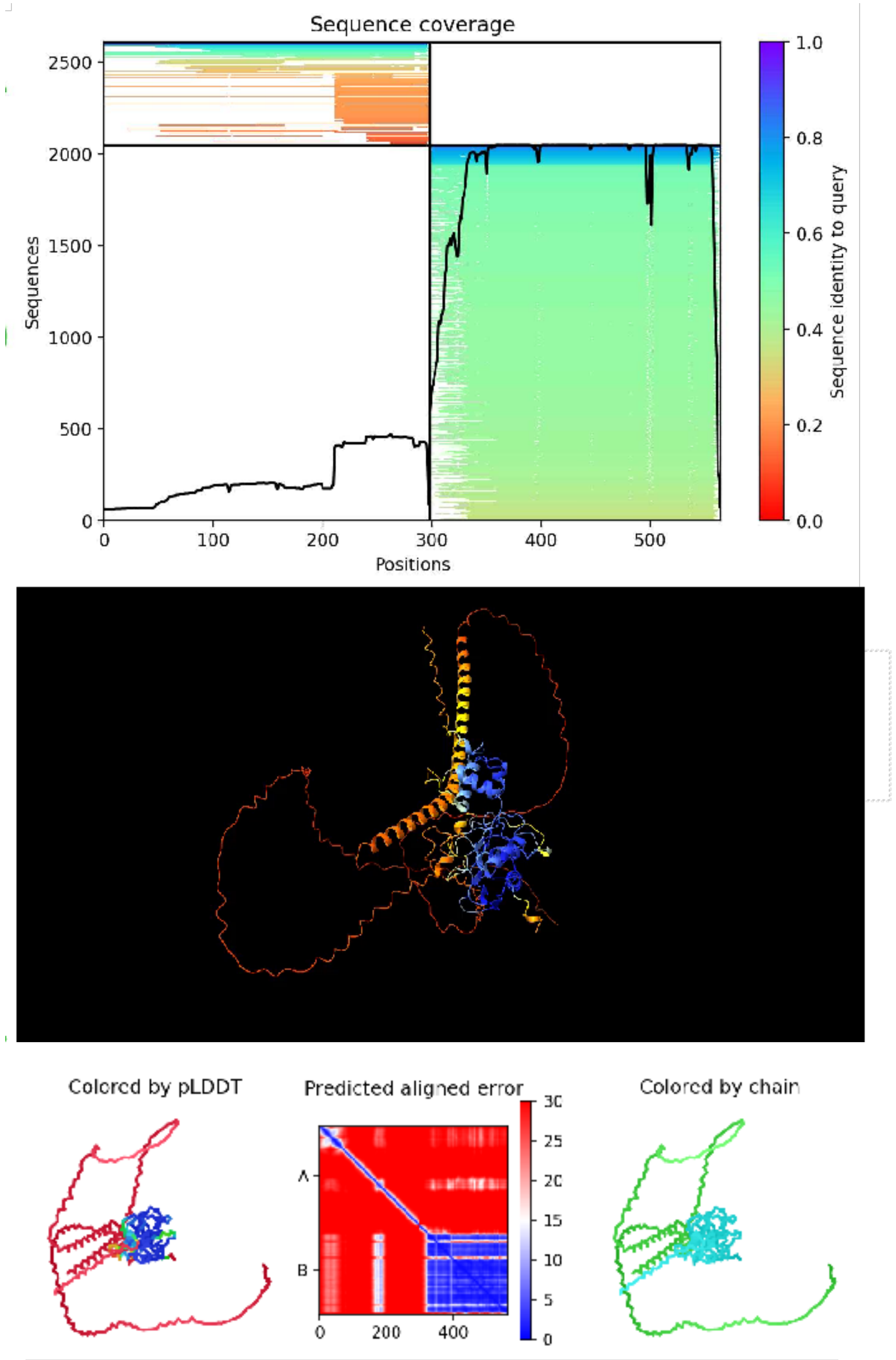


MMP7-G Predicted Docking

B

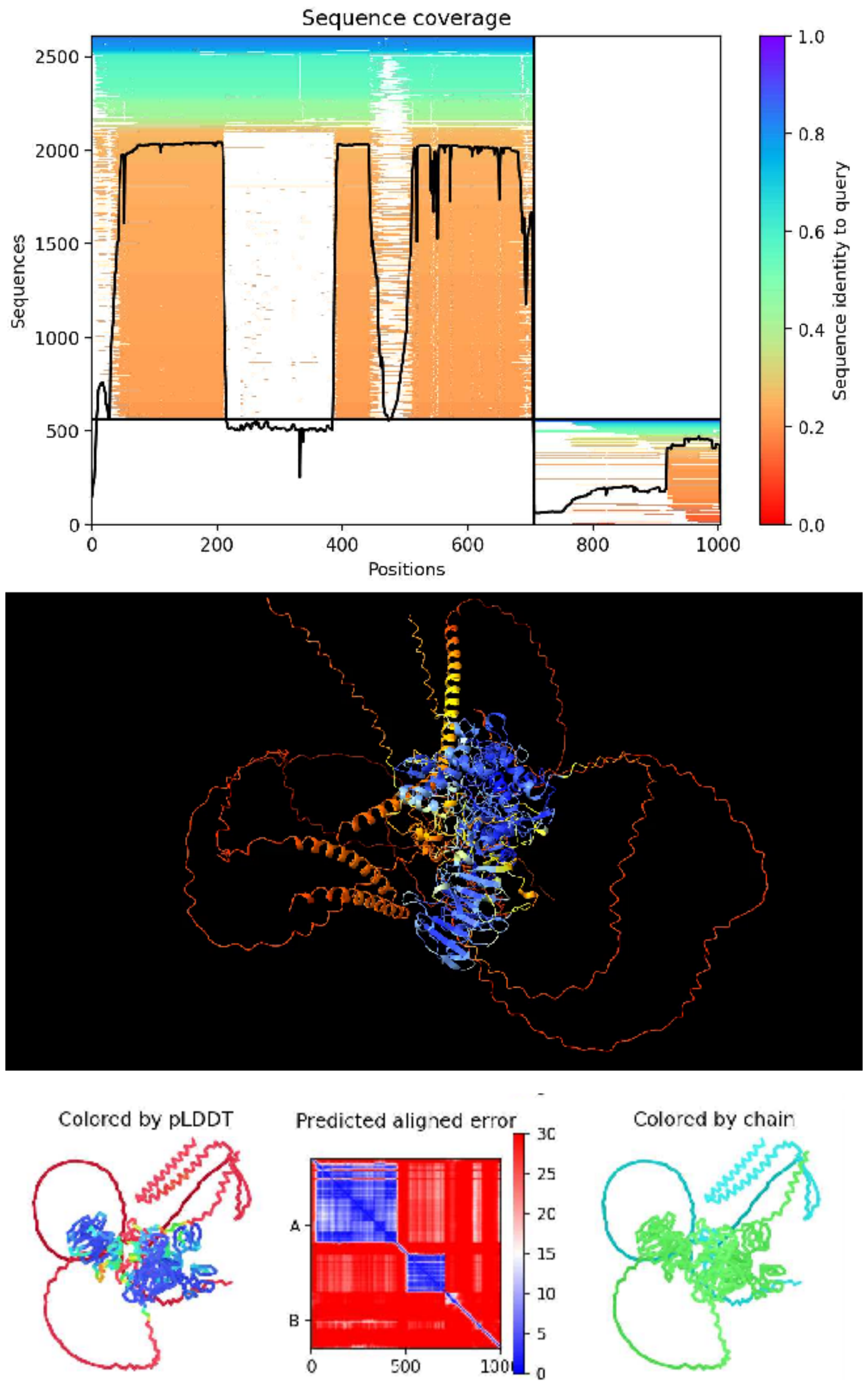


MMP9-G Predicted Docking

**Fig. S3**

**Supplemental information**

**A**

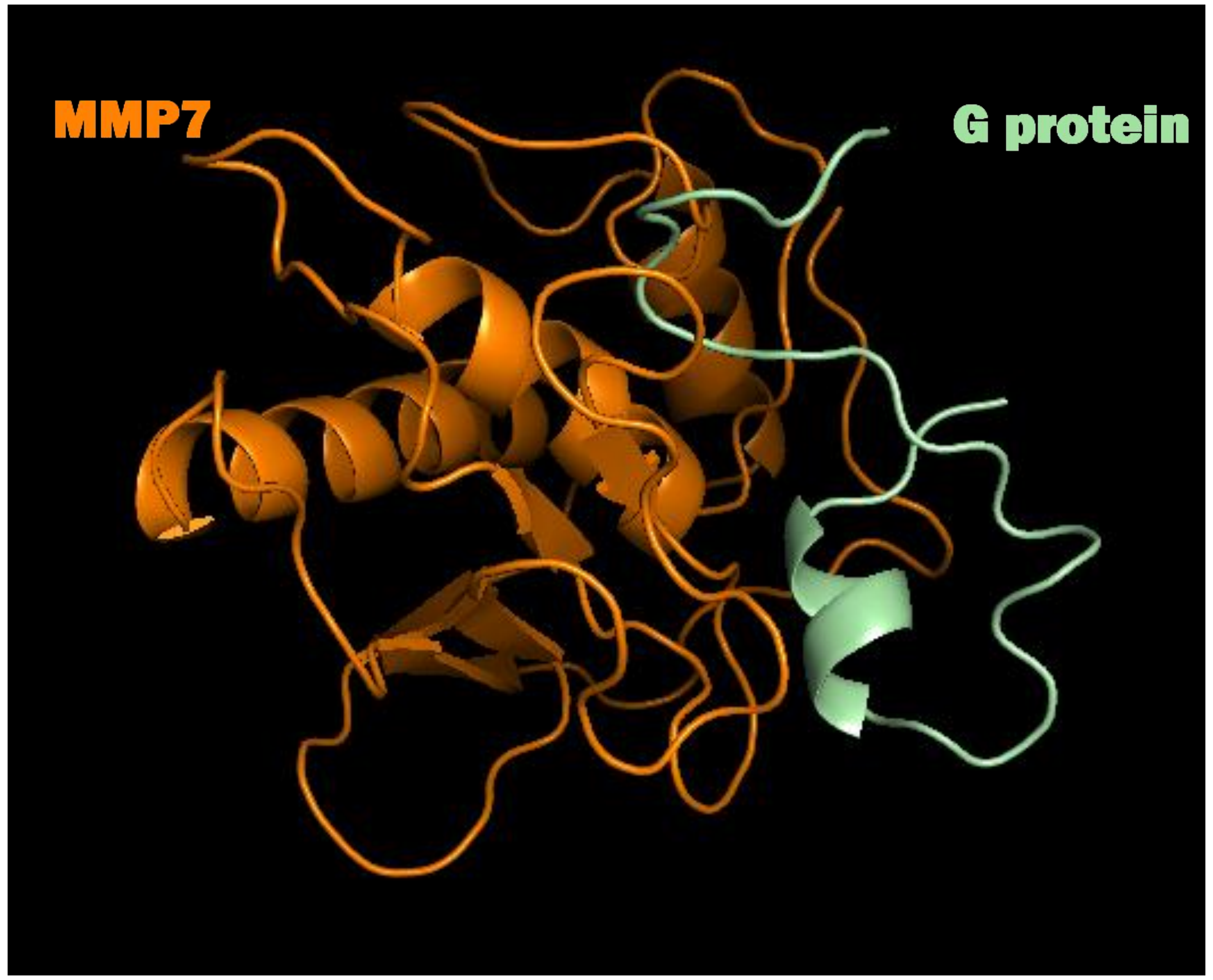


G-MMP7 Docking

**B**

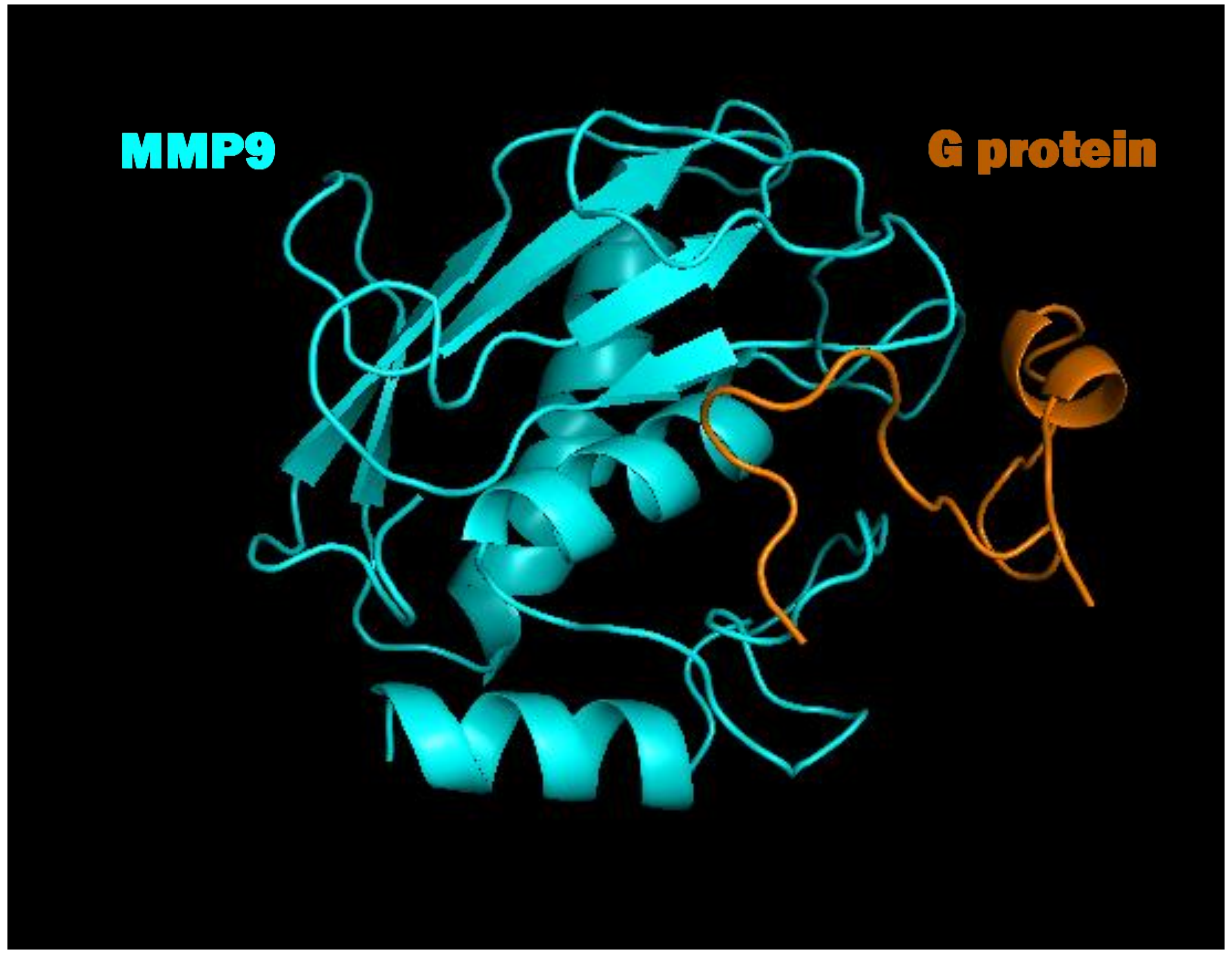


G-MMP9 Docking

**Fig. S4**

**Supplemental information**

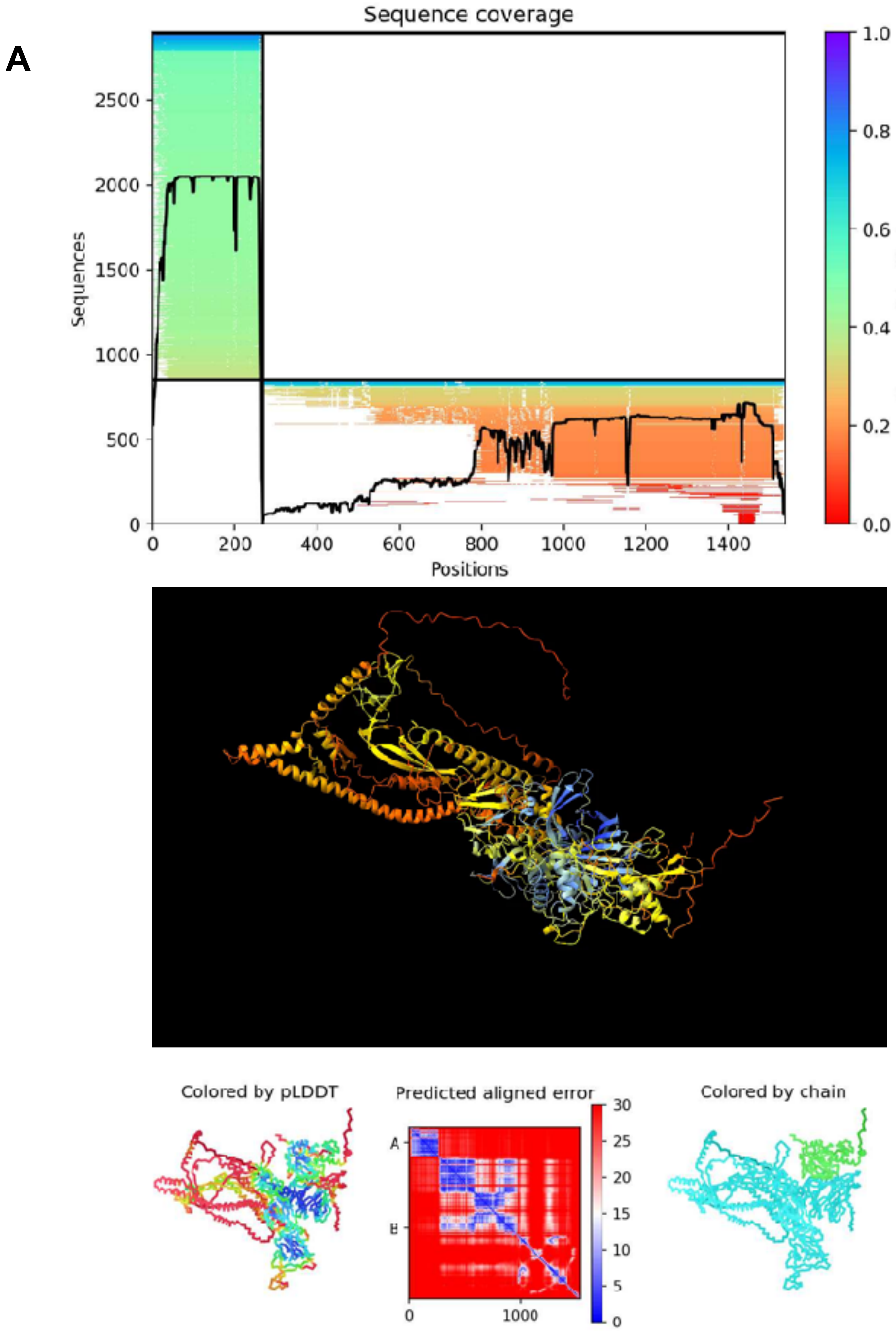


Spike Protein-MMP7 Predicted Docking

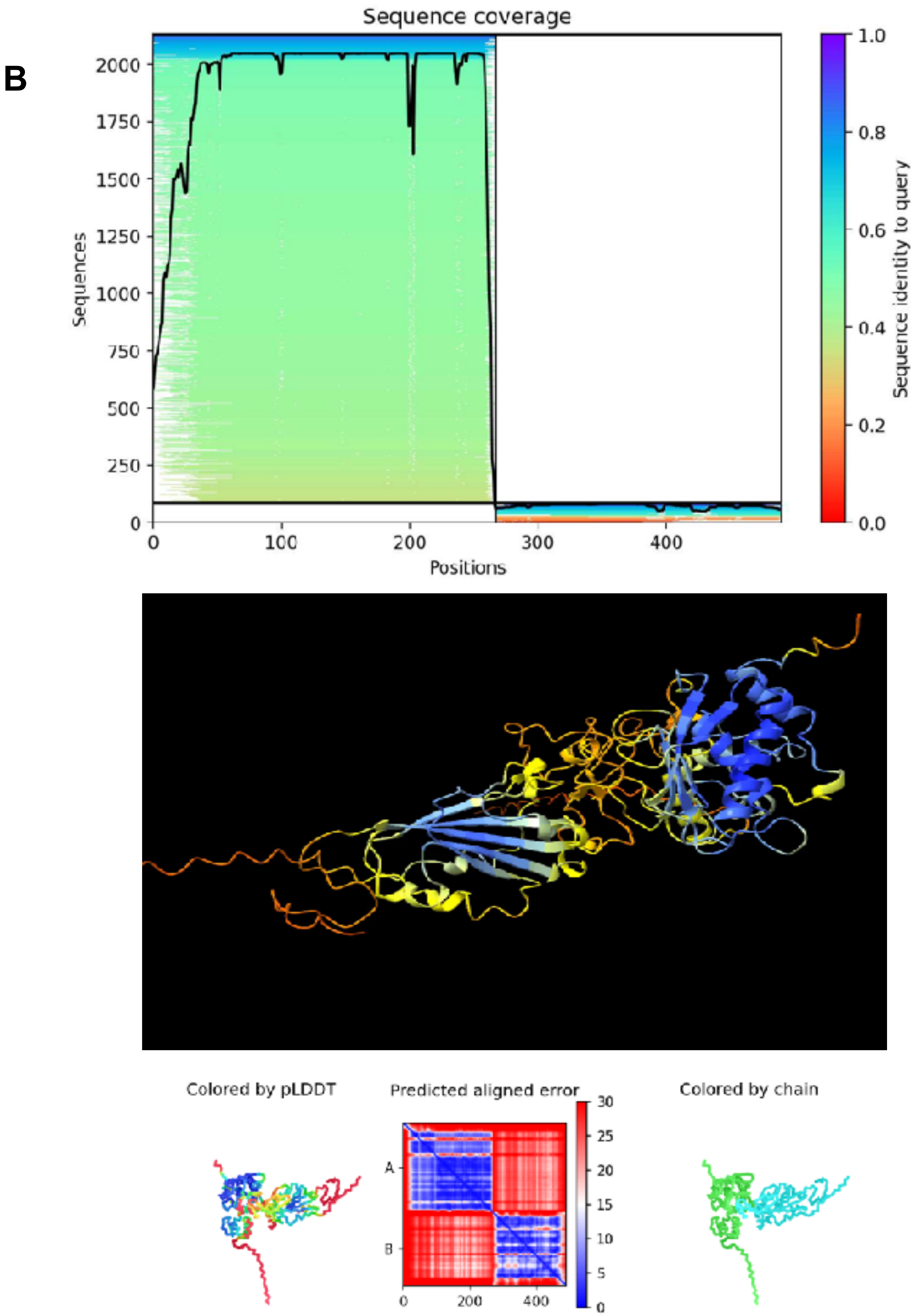


RBD -MMP7 Predicted Docking

**<u>Fig. S5</u>** **Supplemental information**

A

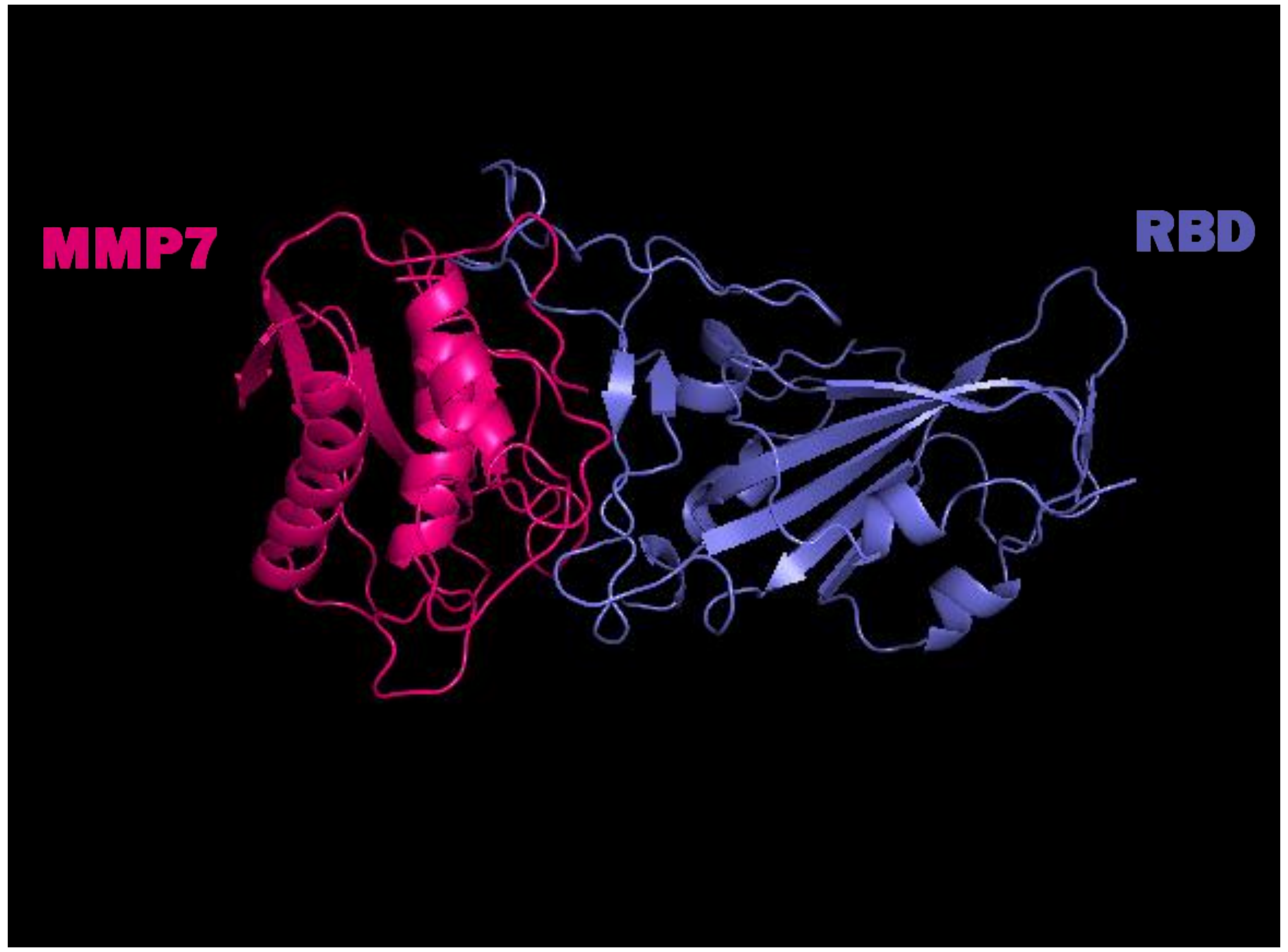


RBD-MMP7 Docking

B

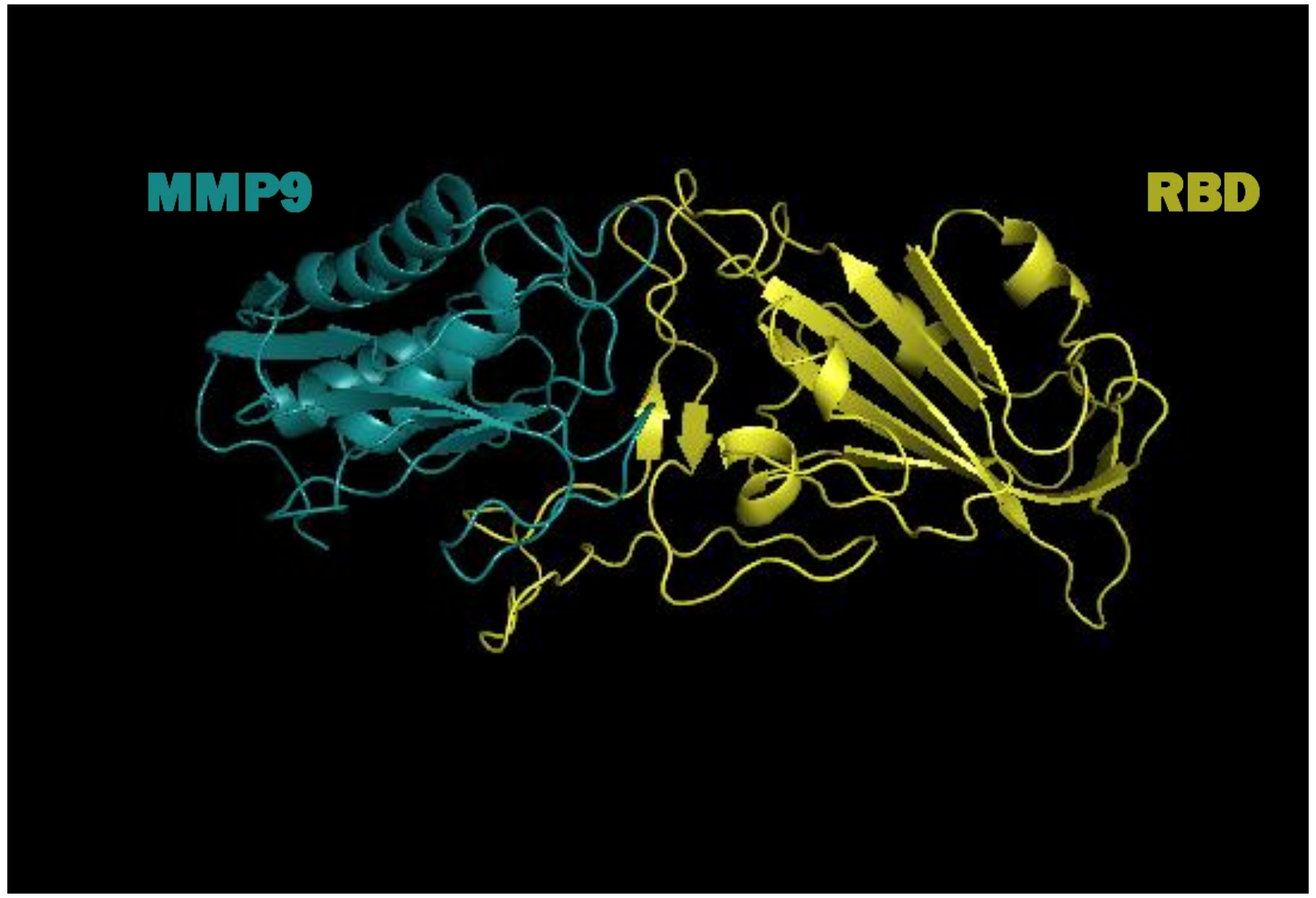


RBD-MMP9 Docking
